# Molecular drivers of insecticide resistance in the Sahelo-Sudanian populations of a major malaria vector *Anopheles coluzzii*

**DOI:** 10.1101/2022.03.21.485146

**Authors:** Sulaiman S. Ibrahim, Abdullahi Muhammad, Jack Hearn, Gareth D. Weedall, Sanjay C. Nagi, Muhammad M. Mukhtar, Amen N. Fadel, Leon J. Mugenzi, Edward I. Patterson, Helen Irving, Charles S. Wondji

## Abstract

Information on common markers of metabolic resistance in malaria vectors from countries sharing similar eco-climatic characteristics can facilitate coordination of malaria control. Here, we characterized populations of the major malaria vector *Anopheles coluzzii* from Sahel region, spanning four sub-Saharan African countries: Nigeria, Niger, Chad and Cameroon. Genome-wide transcriptional analysis identified major genes previously implicated in pyrethroid and/or cross resistance to other insecticides, overexpressed across the Sahel, including CYP450s, glutathione S-transferases, carboxylesterases, and cuticular proteins. Several, well-known markers of insecticide resistance were found in high frequencies - including in the voltage-gated sodium channel (V402L, I940T, L995F, I1527T and N1570Y), the *acetylcholinesterase*-1 gene (G280S) and the *CYP4J5*-L43F (fixed). High frequencies of the epidemiologically important chromosomal inversions, 2La, 2Rb and 2Rc were observed (∼80% for 2Rb and 2Rc). The 2La alternative arrangement is fixed across the Sahel. Low frequencies of these inversions (<10%) were observed in the fully insecticide susceptible laboratory colony of *An. coluzzii* (Ngoussou). Several of the most commonly overexpressed metabolic resistance genes sit in these three inversions. Two commonly overexpressed genes, *GSTe2* and *CYP6Z2* were functionally validated. Transgenic *Drosophila melanogaster* expressing *GSTe2* exhibited extremely high DDT and permethrin resistance (mortalities < 10% in 24 h). Serial deletion of the 5’ intergenic region, to identify putative nucleotide(s) associated with *GSTe2* overexpression, revealed that simultaneous insertion of adenine nucleotide and a transition (T->C), between Fork-head box L1 and c-EST putative binding sites were responsible for the high overexpression of *GSTe2* in the resistant mosquitoes. Transgenic flies expressing *CYP6Z2* exhibited marginal resistance towards 3-phenoxybenzylalcohol (a primary product of pyrethroid hydrolysis by carboxylesterases) and a type II pyrethroid, α-cypermethrin. However, significantly higher mortalities were observed in *CYP6Z2* transgenic flies compared with controls, on exposure to the neonicotinoid, clothianidin. This suggests a possible bioactivation of clothianidin into a toxic intermediate, which if true make it an ideal insecticide against populations of *An. coluzzii* overexpressing this P450. These findings will facilitate regional collaborations within the Sahel region, and refine implementation strategies through re-focusing interventions, improving evidence-based, cross-border policy towards local and regional malaria pre-elimination.

## Background

Since the year 2000 the massive scale-up in vector control interventions and treatment with antimalarial drugs had cut malaria incidence by ∼40 % across Africa (1). Bolstered by this progress the World Health Organization (WHO) has been pushing to eliminate malaria, as proposed in the Global Technical Strategy (GTS) 2016-2030 (2). Unfortunately, the GTS, an ambitious framework with targets to reduce global malaria burden by 90 % in 15 years was dealt an immediate blow by a rebound in malaria transmission, with increased cases between 2016 and 2019 (3, 4). This stark warning of the risk posed to control and elimination efforts was a reflection of the lack of progress in the primary regions of interest in sub-Saharan Africa, which constitute 96 % of the 627,000 malaria-related deaths in 2020 alone (4, 5). Indeed, as the WHO widens the net of malaria elimination (E-2025) its important to acknowledge that no meaningful progress will be made without progress in sub-Saharan Africa, where none of the six countries having the highest global burden of malaria (e.g. Nigeria alone contributing ∼27 % of all cases) is on the path to elimination (6).

The recent escalation of insecticide resistance in the major malaria vectors (7–9) makes the development of molecular tools to anticipate emergence and predict spread of resistance across the African continent imperative, in order to achieve malaria control and elimination. For decades no simple molecular assays were available to track metabolic resistance and assess its impact on malaria control and transmission, until the recent discovery of two DNA markers in the *cis*-regulatory region of two cytochrome P450s, *CYP6P9a* (10) and *CYP6P9b* (11) in the major malaria vector *Anopheles funestus*; findings which allowed the design of simple PCR assays to detect and track metabolic resistance in the field. Unfortunately, these markers explain resistance only in southern Africa for *An. funestus*, since genetic basis of pyrethroid resistance, and cross-resistance with other insecticides is complex in other African regions with different fronts, driven by distinct genes in the presence of barriers to gene flow (10, 12, 13). No major metabolic resistance markers for P450s or GSTs (functionally validated and characterised for epidemiological impact in the natural populations) exist for the major malaria vectors of the *Anopheles gambiae* Complex, though it is the omnipresent vector species with widespread presence across Africa (14). This is hindering early detection and tracking of molecular drivers of resistance in this species, slowing down evidence-based control measures and resistance management.

Major genomic regions associated with metabolic resistance to pyrethroids in *An. gambiae* sensu lato include the *CYP6* P450 clusters on the 2R and 3R chromosomes, and a *GST* epsilon (*GSTe*) cluster on chromosome 3R (15), with some key resistance-associated genes functionally validated. These genes include *CYP6P3* shown to confer cross resistance to pyrethroids and organophosphates (16, 17), *CYP6M2* conferring cross resistance to pyrethroids (18, 19) and DDT (19), as well as *GSTe2* shown to confer resistance to DDT (20). For *GSTe2*, in addition to the I114T marker (20), a recent study has found a novel mutation (*Gste2*-119V) associated with resistance (21) using a high-throughput genotypic panel for markers. Functional validation and fild data is required establish the empirical evidence of the role of this mutation in resistance. What has been missing in the case of *Anopheles gambiae* and *An. coluzzii* is reliable molecular markers of resistance for major metabolic gene families, e.g. P450s (with field and laboratory validated data) to aid creation of DNA-based diagnostic assays which will allow (i) easy tracking of resistance in the field (e.g. the case of 119F-*GSTe2* mutation in *An. funestus* (22), and (ii) determination of the operational impact of the resistance markers in the field, as recently done for 119F-*GSTe2* (23), *CYP6P9a*_R and *CYP6P9b*_R markers (10, 11) in *An. funestus*.

The *An. gambiae s.l.*, especially populations in the semi-arid steppe, exhibit high frequency of paracentric chromosomal inversions [one of the most effective instruments for speciation and local adaptations (Ayala et al., 2014, Dobzhansky, 1971, Kirkpatrick, 2010)], maintained in spatially and temporally heterogenous environment, and which segregate along climatic gradients of increasing aridity (24)]. The 2La inversions are associated with resistance to desiccation in adults (Fouet et al., 2012, Gray et al., 2009) and thermal stress in larvae (Cassone et al., 2011). It was also shown that inversion 2La assorts with insecticide resistance, e.g., dieldrin plus fipronil (Brooke et al., 2000), and is associated with thermotolerance and permethrin resistance in the Sahelain *An. coluzzii* (Ibrahim et al., 2021).

To support malaria pre-elimination effort in sub-Saharan Africa we targeted the Sahelo-Sudanian region, which represent northern-most limit of malaria endemicity in sub-Saharan Africa, and where malaria is highly seasonal [offering excellent target for pre-elimination effort through sustained seasonal vector control and seasonal malaria chemoprevention (25)]. Focusing on the predominant malaria vector, *An. coluzzii* from Sudano-Sahelian transects of four countries, Nigeria, Niger, Chad and Cameroon (7, 26–28) we identified the major metabolic resistance genes mediating pyrethroid resistance and cross-resistance in this region, established the genetic variants which explained the resistance. We also functionally validated the roles of two major candidate genes in the resistance (*CYP6Z2* and *GSTe2*) using transgenic *Drosophila melanogaster* flies (GAL4/UAS system), as well as identifying single nucleotide polymorphisms in the 5’ regulatory elements of *GSTe2,* responsible for its overexpression in the resistant population.

## Materials and Methods

### Study Site and Mosquito Sampling

Blood fed female *An. coluzzii* mosquitoes, resting indoor were collected at one locality each in sub-Sahel (Additional Figure S1): Hadiyau (HAD: 12°21′38″N, 9°59′15″E), a sub-Sahel village in northern Nigeria; Takatsaba (TAK: 13°44’01.8"N 7°59’05.2"E), a Sahel village in southern Niger; Simatou (SIMAT: 10°50’40.7"N 14°56’40.9"E), a sub-Sahel village in Maga Department, far north of Cameroon; and Massakory (CHAD: 12° 6′ N, 15° 02′ E), a Sahel town in Chad Republic. Details of sampling approaches and resistance profiles of mosquitoes collected from Nigeria, Niger, and Chad are available in previously published articles (7, 26, 28). As in the above countries the Simatou F_1_ females were also highly pyrethroid resistant, with a mortality of only 3.7% from WHO tube bioassays using 0.05% deltamethrin, and no mortality at all with 0.75% permethrin (data not published).

### Genome-wide transcriptional analysis for common insecticide resistance genes in the Sahel regions

#### RNA extraction, library preparation and sequencing

The RNA was extracted using the Arcturus PicoPure RNA isolation Kit (Applied Biosystems, CA, USA) from three pools of 8 F_1_ *An. coluzzii* females (2-4 day old from the same population) alive after exposure to deltamethrin (resistant, R), unexposed (control, C), and also from unexposed females of the fully susceptible laboratory colony of *An. coluzzii*, Ngoussou (susceptible, S) (29). The RNA isolation was carried out following the manufacturer’s protocol with *Dnase* I-treatment to remove contaminating DNA. The quantity and quality of RNA was measured using a NanoDrop spectrophotometer (ThermoFisher, MA, USA) and Bioanalyzer (Agilent, CA, USA).

Library preparation, sequencing and data quality control were carried out by the Centre for Genomic Research (CGR), University of Liverpool, UK. RNA samples were subjected to poly(A) mRNA enrichment and libraries prepared from the poly(A) mRNA-enriched materials (dual-indexed, strand-specific RNAseq libraries were prepared using the NEBNext polyA selection and Ultra Directional RNA library preparation kits). Libraries were sequenced on a single lane of an Illumina HiSeq 4000 (paired-end, 2x150 bp sequencing, generating data from >280 M clusters per lane). Basecalling and de-multiplexing of indexed reads were performed by CASAVA version 1.8.2 (Illumina). De-multiplexed fastq files were trimmed to remove Illumina adapter sequences using Cutadapt version 1.2.1 (30). Option -O 3 was used, so that the 3’ end of any reads which matched the adapter sequence for 3 bp or more were trimmed. Reads were further trimmed to remove low quality bases using Sickle version 1.200 (31) with a minimum window quality score of 20. Reads shorter than 20 bp after trimming were removed. If both reads from a pair passed this filter, each was included in either the R1 (forward reads) or R2 (reverse reads) file. If only one of a read pair passed this filter, it is included in the R0 (unpaired) file. Statistics were generated using fastq-stats from EAUtils (32). Summary of total number of reads for each sample and distribution of trimmed read length for forward (R1) and reverse (R2) reads and reads unpaired after trimming (R0) are provided in Additional Figure S2.

#### Data analysis and estimation of transcript abundance by tag counting and differential gene expression

Paired data for each replicate per country was aligned to the *An. gambiae* reference transcript ome AgamP4.10 downloaded from VectorBase (https://vectorbase.org/) in salmon (0.11.4), using ‘validate mappings’, ‘seqBias’, ‘gcBias’ and ‘rangeFactorizationBins 4’ flags. Read mapping results (pre-alignment and post-alignment descriptive statistics (flagstat output files) showing sequencing depth and coverage are given in Additional Tables 1 and -2, respectively. Salmon results were converted into a gene expression matrix using the Bioconductor package ‘tximport’ for input to DESeq2 1.26.0 (33). Differential gene expression was tested for the three possible combinations of Exposed (R, deltamethrin resistant), Unexposed (C, control) and Susceptible replicates (S). For results interpretation log_2_-fold change thresholds of 1 was imposed, with false discovery rate adjusted p-values of 0.05 applied to accept significance. Principal components analysis implemented in DESeq2 was used to examine relationships between respective replicates and treatments. This was carried out based on the 500 most variable genes, with data transformation (normalisation/scaling) as implemented in VST (DESeq2). For visualisation of expression results, volcano plots were created using the Enhanced Volcano package (34) using the top most overexpressed genes from lists which were prepared with log_2_FC cut off of 1 and p value of 0.01. Heatmaps were generated (35) using the list of the top 50 most overexpressed metabolic resistance genes with log_2_FC cut off of 1 and p value of 0.01.

#### Gene Ontology Enrichment and Mapping/Functional annotations

The enrichment analysis for GO terms was carried out using a topGO package (36) against *An. gambiae* (AgamP4.10), with data reannotated using EggNOG v5.0 (http://eggnog5.embl.de/#/app/home). Gene lists used in topGO are from the log_2_FC = 0, p <0.05 analysis (the default in DESeq2). For the data from each site (country) six set of results were generated (R, C and S contrasts) for genes either up- or down-regulated in each contrast, for molecular function (MF) and biological process (BP) ontologies. For analysis the GO terms lists for MF were used as input into the Revigo (37) for interpretation and visualisation, querying the Whole Uniprot database, and SimRel setting for semantic similarity measurements.

### Quantitative PCR measurement of expression profiles of metabolic resistance genes

The level of expression of 12 resistance-associated genes was validated by qRT-PCR, using the primers provided in Additional Table 3. These include the *GSTe2* (AGAP009194), *GSTZ1* (AGAP002898), *CYP6Z2* (AGAP008218), *CYP6Z3* (AGAP008217), *CYP4C27* (AGAP009246), *CYP4G16* (AGAP001076), *CYP4G17* (AGAP000877), *CYP6P3* (AGAP002865), *CYP6M2* (AGAP008212), *CYP9K1* (AGAP000818), a *UGT-B19* (AGAP007920) and *COEBE3C* (AGAP005372). The qRT-PCR was carried out using three technical replicates each of cDNA extracted from 1 µg of total RNA of three biological replicates each from the Resistant (R), Control (C) and Ngoussou (S). Protocol followed was as established in previous studies (38), with relative expression level and fold change (FC) of each target gene in R and C relative to S calculated according to the 2^-ΔΔCT^ method incorporating the PCR efficiency (39), after normalization with the housekeeping genes ribosomal protein S7, *RPS7* (AGAP010592) and glycerol-3-phosphate dehydrogenase, *GPDH* (AGAP007593). Significant differences were calculated using ANOVA with Dunnett’s post hoc test.

#### Detection of Signatures of Selective Sweep

To detect signature of selective sweeps in the major metabolic resistance genes of interest, a Snakemake RNAseq population genetics pipeline, the RNAseq-Pop was utilised (40). The workflow aligns RNA-Seq reads to the reference genome, and calls genomic variants with *Freebayes*, at a user-provided level of ploidy, in our case 16 (8 diploid pooled mosquitoes). F_st_ (41) per gene between population pairs and Tajima’s D per gene within population were estimated. This was performed against all SNPs passing quality and missingness filters. Population branch statistic (PBS) scans was performed with the Snakemake, conditional on the presence of three suitable populations (42). Hudson’s F_st_ and PBS scans were ran, taking the average for each protein-coding gene, as opposed to in windows. The population genetics statistical analyses were calculated in scikit-allel v1.2.1 (43).

#### Establishment of allele frequencies of variants in genes of interest

After genome alignment, RNA-Seq-Pop utilises samtools (44) to query specific positions of the genome, calculating raw allele frequencies at those sites with a custom R script.

#### Detection of chromosomal inversion polymorphisms and metabolic genes within its breakpoint

A modified version of the Python 3 program, compkaryo (45) was used to karyoptype the major, *An. coluzzii*/*gambiae* phenotypically important inversion polymorphisms in chromosome 2, and calculate its frequencies, *in silico*, using the previously identified tag SNPs significantly associated with inversions. This allows to predict with high confidence genotypes of the six common polymorphic inversions on chromosome 2, in the sequenced field *An. coluzzii,* as well as in the Ngoussou. Compkaryo uses the Ag1000 database (The *Anopheles gambiae* 1000 Genomes Consortium 2017) (15) by leveraging a subset of cytologically karyotyped specimens to develop a computational approach for karyotyping applicable to whole genome sequence. Modifications in the Snakemake pipeline allows for variable ploidy (useful in the case of replicates from our pooled RNAsequencing samples) here.

### Functional validation of the commonly overexpressed resistance-associated genes

#### Comparative analysis of coding sequences of major resistance genes

Two resistance-associated genes, *GSTe2* and *CYP6Z2* which feature most premoninently in the resistant *An. coluzzii* populations across the Sahel were chosen for functional validation. To establish presence of allelic variants which could impact catalytic activities, full length coding sequences (cDNAs) of *GSTe2* and *CYP6Z2* were amplified and sequenced from alive mosquitoes in the four Sahel countries, as well as from the Ngoussou. This was done using total RNA extracted from 5 individual pools of 8 F_1_ *An. coluzzii* females (2-4 day old) alive after exposure to deltamethrin (resistant, R for *CYP6Z2*) or DDT (R for *GSTe2*). Protocol for RNA extraction was as described in previous section, above. Amplification was done using Phusion HotStart II Taq Polymerase (ThermoFisher SCIENTIFIC, MA, USA) and the Full primers listed in Additional Table 4. The PCR mix comprised 5x Phusion HF Buffer (containing 1.5 mM MgCl_2_), 85.7 µM deoxynucleotides (dNTPs), 0.34 µM each of forward and reverse primers, 0.015 U of Phusion HotStart II DNA Polymerase (Fermentas, MA, USA), 10.71 µL of ddH_2_0 and 1 µL cDNA. Thermocycling conditions were 1 cycle at 95 °C for 5 min; followed by 35 cycles each of 94 °C for 20 s, 60 °C for 30 s, 72 °C for 2 min (1 min for *GSTe2*); and finally, one cycle at 72 °C for 5 min. PCR products were cleaned with a QIAquick^®^ PCR Purification Kit (QIAGEN, Hilden, Germany) and ligated into the pJET1.2/blunt cloning vector using the CloneJET PCR Cloning Kit (ThermoFisher SCIENTIFIC, MA, USA). These were then cloned into *E. coli DH5α,* plasmids miniprepped with the QIAprep^®^ Spin Miniprep Kit (QIAGEN) and sequenced on both strands using pJET1.2 primers.

Polymorphisms were detected through examination and manual editing of sequence traces using BioEdit version 7.2.3.0 (46) and nucleotide differences in sequences aligned using CLC Sequence Viewer 7.0 (http://www.clcbio.com/). Different haplotypes were compared by constructing a maximum likelihood phylogenetic tree using MEGA X (47). Genetic parameters of polymorphism including number of haplotypes (h) and its diversity (H_d_), number of polymorphic sites (S) and nucleotide diversity (π) were computed using DnaSP v6.12.03 (48).

#### Characterization of the 5’ regulatory regions of GSTe2 and CYP6Z2

##### Amplification, cloning and sequence characterisation of 5’ regulatory element

To investigate presence of genetic variants in the regulatory elements, which could be responsible for overexpression of *GSTe2*, 351 bp intergenic regions (spanning the 43 bp 3’ UTR of *GSTe1,* 248 bp flanking sequence and 60 bp 5’UTR of *GSTe2*) preceding the start codon were amplified from 10 each of DDT-alive and -dead females from the 4 Sahel countries, as well as from the Ngoussou females (primers provided in Additional Table 4). For *CYP6Z2,* a 1078 bp intergenic region was retrieved from the VectorBase and used for amplification of the putative 5’-regulatory elements. Primers spanning 38 bp 3’ UTR of *CYP6Z1,* a 937 bp flanking sequence and 103 bp 5’UTR of *CYP6Z2,* preceding the start codon of *CYP6Z2* were used to amplify fragments from 10 each of deltamethrin-alive and -dead females from Nigeria and Niger, as well as from the Ngoussou females. Amplification was carried out using HotStart II Polymerase (ThermoFisher SCIENTIFIC, MA, USA) with similar thermocycling conditions as outlined above for coding region of *GSTe2* and *CYP6Z2,* respectively. Purification of PCR amplicons, cloning into pJET1.2 vector, sequencing and polymorphism analysis were done as outlined above.

The 351 bp 5’-UTR fragments of *GSTe2* and 1078 bp fragment of the *CYP6Z2* were analysed with the Gene Promoter Miner (http://gpminer.mbc.nctu.edu.tw/) and MatInspector (49) to identify putative promoter elements, predict transcription start (TSS) and potential transcription factor binding sites.

##### Cloning of GSTe2 and CYP6Z2 5’ regulatory elements in PGL3-Basic vector and dual luciferase reporter assay

Following analysis of the above sequences the 351 bp intergenic fragments of *GSTe2* were amplified from the most predominant sequences of DDT-alive, DDT-dead and Ngoussou. Same was done for for *CYP6Z2* amplifying 1087 bp fragment from deltamethrin-alive and - dead, and Ngoussou. Primers bearing *kpn*I and *BgL*II sites (Additional Table 4) allowed incorporation into pGL3-Basic reporter vector containing luciferase gene from the firefly *Photinus pyralis* (Promega, Wisconsin, USA). Amplification was carried out using Phusion HotStart II Polymerase, with conditions as above, followed by purification of PCR amplicons, and cloning into pJET1.2 vector. Positive colonies (sequencing primers for pGL3-Basic provided in Additional Table 4) were miniprepped; the minipreps digested with the above restriction enzymes, gel-purified and ligated upstream of luciferase gene in pGL3-Basic vector already linearized with the same restriction enzymes. Positive colonies were cloned and miniprepped and concentrations of the recombinant plasmids adjusted to 200 ng/µL before co-transfection. Protocols for maintenance of the *An. gambiae* cell line 4a-3B (MRA-919, https://www.beiresources.org/), co-transfection of 200 ng of either *GSTe2* or *CYP6Z2* promoter constructs, LRIM promoter in pGL3-Basic vector (50), or promoter-less pGL3-Basic control, with 1 ng/µL of internal control, sea pansy *Renilla reniformis* luciferase containing the *Drosophila* Actin *5C* promoter in pRL-null (50), are provided in Additional File 1 (additional text for Methods and Results). Protocol for cell lysis and measurement of the activities of the firefly luciferase (Dual-Luciferase Reporter Assay kit, Promega), with normalisation using the Renilla luciferase activity, were as outlined in the Promega Quick Protocol, and provided in detail in Additional File 1.

Assay background was also measured using lysate from non-transfected control cells. Results were compared using a two-tailed Chi-Square test of independence using GraphPad Prism 7.02 (GraphPad Inc., La Jolla, CA, USA).

##### Generation of GSTe2 promoter deletion constructs and assays

The intergenic region separating *GSTe1* from *GSTe2* was progressively delineated. Partial fragments of the 5’ regions (encompassing the 3’-UTR of *GSTe1*, the *GSTe1*/*GSTe2* flanking region and the 5’-UTR of the *GSTe2*) were created by sequential deletion using forward primers (provided in Additional Table 4, with numbers in primer names referring to distances from the AUG start codon of *GSTe2*). Reverse primers were those initially used for amplification of the full 351 bp nucleotide fragments. These fragments include (i) 308 nucleotides fragment generated from deletion of the 43 bp 3’-UTR of the *GSTe1* (-308 primers) obliterating cellular-Myb (c-Myb) transcriptional factor binding site; (ii) 291 nucleotide fragments generated from deletion of *GSTe1* 3’-UTR plus 38 nucleotides of the flanking region (-270 primers), which removed in addition *δ* elongation factor 1 (*δ*EF1) binding site; (iii) 262 nucleotide fragments produced from deletion of *GSTe1* 3’-UTR plus 46 nucleotides of the flanking region (-262 primers), which obliterated in addition Forkhead box L1 (FOX-L1) putative binding site; as well as (iv) 232 nucleotide fragments produced from deletion of *GSTe1* 3’-UTR plus 75 nucleotides of the flanking region (-232 primers), located 11 nucleotides from the c-EST/grainy head transcription factor binding site. Also, to assess the importance of the *GSTe2* 5’UTR, 43 nucleotides upstream the AUG codon were deleted (from the 60 nucleotide 5’-UTR of both the Sahel-alive and Ngoussou), leaving the putative transcription start site motif untouched. This was done using the original forward primer for the 351 nucleotides intergenic region, with a newly designed primer, GSTe2_minus_5’-UTR-R (Additional Table 4).

#### Characterisation of major resistance genes using transgenic analysis

##### Cloning and microinjection of GSTe2 and CYP6Z2 into Drosophila melanogaster

Transgenic flies expressing recombinant GSTe2 and CYP6Z2 were created using GAL4/UAS system, as outlined in a previous publication (51) and used in contact bioassays, to confirm if over-expression of these genes alone can confer resistance to insecticides. Details of amplification of full-length *GSTe2* and *CYP6Z2* from the predominant alleles of sequences of cDNA, cloning into pUASattB vector, microinjection into germ-line *D. melanogaster* lines, crossing of the UAS-lines with the GAL4-Actin driver strain are outlined in Additional File 1.

To confirm overexpression of *GSTe2* and *CYP6Z2* in the transgenic flies three replicates each of 6 females (both experimental and control group) were used for qRT-PCR using a previously established protocol (38). Total RNA and cDNA were extracted as described above, and the relative expression levels of transgenes were assessed, with normalization using the *RPL11* housekeeping gene. The qtrg primers used for the two genes and the *RPL11* primers are provided in Additional Table 4.

##### Insecticide Susceptibility Contact Bioassay

For insecticide bioassays, 3- to 4-day old experimental and control F_1_ females were exposed to 0.15 % deltamethrin, 2 % permethrin, 0.05 % α-cypermethrin, 4 % DDT and 2 % clothianidin-impregnated papers prepared in acetone and Dow Corning 556 Silicone Fluid (BHD/Merck, Hesse, Germany). Flies overexpressing *CYP6Z2* were also exposed to the primary product of hydrolysis of pyrethroids: 4 % and 20 % (5x) 3-phenoxybenzaldehyde (PBAld) and 3-phenoxybenzylalcohol (PBAlc). Transgenic flies expressing *GSTe2* were exposed to 2 % permethrin, 0.15 % deltamethrin, 0.05 % α-cypermethrin and 4 % DDT only. Impregnated papers were rolled and introduced into 45 cc plastic vials to cover the entire wall and the vials plugged with cotton soaked in 10 % sucrose (38). 20–25 flies were placed in each vial, and the mortality plus knockdown scored at 1 h, 3 h, 6 h, 12 h and 24 h of exposure to the insecticides. For each insecticide, assays were performed in 6 replicates and Student’s t-test used to compare the mortality plus knockdown between the experimental groups and the control. Controls were adult F_1_ progeny with the same genetic background as the experimental group but without the *GSTe2* or *CYP6Z2* inserts. They were obtained by crossing virgin females from the driver strain Act5C-GAL4 and the UAS recipient male lines with white eyes (not carrying the pUASattB-GSTe2 or pUASattB-CYP6Z2 insertions).

## Results

### Genome-wide transcriptional profile of the Sahelian *An. coluzzii* populations

A three-way pairwise comparison was conducted for the data from each country: resistant *vs* susceptible (R-S), resistant *vs* unexposed control (R-C) and unexposed control *vs* susceptible (C-S). This captures background variations due to geographical differences in the resistant vs susceptible (R-S) comparison, accounts for genes overexpressed due to induction (R-C comparison), as well as genes that are constitutively overexpressed (C-S comparison). A total of 1384 genes were significantly differentially expressed (FDR-adjusted p <0.05 and log_2_ fold change threshold of 1/FC ≥ 2) in R-S comparison in Nigeria (1077 upregulated and 307 down-regulated); 1185 genes were differentially expressed in C-S (1002 upregulated and 183 downregulated); and 295 genes in R-C (129 upregulated and 166 downregulated). Of these, 52 genes were commonly differentially expressed in all 3 comparisons (Figure S3, panel a), including the upregulated genes, *COEAE80* (AGAP006700), *CYP4H18* (AGAP028019), *CYP4H17* (AGAP008358) and cuticular proteins, *CPLCX3* (AGAP006149), *CPR59* (AGAP006829), *CPR76* (AGAP009874) and *CPR75* (AGAP009871). The Additional Figure S3, panels a-d depicts the differentially expressed genes for the four countries. For Niger, 881 genes were differentially expressed in R-S comparison (619 upregulated and 262 down-regulated) (Additional Figure S3b); 1256 genes were differentially expressed in C-S (986 upregulated and 270 downregulated); and 196 genes in R-C (81 upregulated and 115 downregulated). Of these, 22 genes were commonly differentially expressed in all 3 comparisons [including the upregulated *aminopeptidase N1* (AGAP012757), *CYP6Z2* (AGAP008218), *chymotrypsin-3* (AGAP006711) and *chymotrypsin-2* (AGAP006710), and an acid trehalase (AGAP008547)]. For Chad, 1392 genes were differentially expressed in R-S comparison (975 upregulated and 417 down-regulated) (Additonal Figure S3c); 1284 in C-S (105 upregulated and 269 downregulated); and 526 genes in R-C (270 upregulated and 256 downregulated). Of these, 97 genes were commonly differentially expressed in all 3 comparisons [including *CYP4C27* (AGAP009246), *SULTD1* (AGAP012672), *aminopeptidase N1*, and diverse cuticular proteins, e.g. chitinase (*Cht24*, AGAP006191, *CPFL1* (AGAP010902), *CPCFC1* (AGAP007980), *CPLCX3*, *CPR24* (AGAP005999), *CPR106* (AGAP006095) and *CPR130* (AGAP000047). Finally, for Cameroon, 376 genes were differentially expressed in R-S comparison (204 upregulated and 172 down-regulated) (Additional Figure S3d); 932 in C-S (778 upregulated and 154 downregulated); and 116 genes in R-C [only 9 upregulated and 107 downregulated (probably due to a single, low quality replicate in the raw data from Cameroon (Additional Figure S2). Not surprising, only 7 genes were commonly differentially expressed in all 3 comparisons. These include the highly upregulated gene, *GSTe2* (AGAP009194) and *chymotrypsin-1* (AGAP006709).

All data analysed together (Additional Figure S3e) revealed no single gene differentially expressed in common; possibly due to the low quality with the Chad unexposed (C) data. Analysis of data from Nigeria, Niger and Chad, revealed a single gene (AGAP000046, transporter major facilitator superfamily) differentially expressed across all countries (Additional Figure S3f). Niger and Chad shared *hexamerin* (AGAP010658), *aminopeptidase N1*, and unknown protein, AGAP0290967; Nigeria and Niger share only a single gene, AGAP003248; while seven genes were common to Nigeria and Chad, including *CPLCX3*.

Principal component analysis for the top 500 most variable genes in all experimental arms revealed data from field samples (R and C) from all four countries clustering closer in PC1 and PC2 axes, away from the data from the susceptible females, Ngoussou (Additional Figure S4).

#### Analysis of the common differentially expressed genes across the Sahel

The most differentially expressed genes were presented in Figure 1, a volcano plot of fold change *vs* significance levels for Nigeria and Niger, and Figure 2, for Chad and Cameroon. Detailed lists of these genes of interest is provided in Additional File 2. Comparisons of genes commonly, upregulated and/or downregulated in R-S/R-C/C-S, from the four countries revealed similar transcriptomic profiles between R and C compared with the S. The most commonly and consistently overexpressed genes across the Sahel (taking account mean expressions) are the chymotrypsins, 3, -2 and -1 (*CHYM3*/AGAP006711 and *CHYM2*/AGAP006710 and *CHYM1*/AGAP006709) (Figures 1 and 2, Additional File 2), the glutathione S-transferase, *GSTe2* (AGAP009194), an aquaporin, *AQP3* (AGAP010326), *CYP6Z2* (AGAP008218), *CYP6Z3* (AGAP008217), *CYP4C27* (AGAP009246), a chitinase, *Cht24* (AGAP006191), a thioester-containing protein-1, *TEP-1* (AGAP010815),), a trehalose 6-phosphate synthase/phosphatase, *TPS*1/2 (AGAP008227), a lipase (AGAP002353), and AGAP012818 (V-type protein ATPase subunit A). More on these genes is provided in the sections, below.

**Figure 1:** A volcano plot of differentially expressed genes showing fold changes and levels of significance in R-S, C-S and R-C comparisons, for Nigeria and Niger populations of *An. coluzzii*. The plot depicts the top upregulated and top downregulated genes for each comparison, with several genes including chymotrypsins, a malate dehudrogenrase, CYP450s, a glutathione S-transferase (*GSTe2*), and a lipase, commonly upregulated in R vs S and C vs S comparisons. R stands for the female, resistant mosquitoes (survivors of 0.05% deltamethrin, 24 h after exposure), C stands for control (mosquitoes from the same population and of the same age not exposed to deltamethrin, and S stands for susceptible (females fromfully insecticide susceptible colony, Ngoussou).

**Figure 2:** A volcano plot of differentially expressed genes showing fold changes and significance in R-S, C-S and R-C comparisons, for Chad and Cameroon populations of *An. coluzzii*. The plot depicts the top upregulated and the top downregulated genes for each comparison with several genes including chymotrypsins, a malate dehudrogenrase, an aquaporn, CYP450s (particulary *CYP6Z2*), a glutathione S-transferase (*GSTe2*) and a lipase, commonly upregulated in R vs S and C vs S comparisons. R stands for the female, resistant mosquitoes (survivors of 0.05% deltamethrin, 24 h after exposure), C stands for control (mosquitoes from the same population and of the same age not exposed to deltamethrin, and S stands for susceptible (females from fully insecticide susceptible colony, Ngoussou).

Other genes commonly overexpressed in data from two or three countries include AGAP005501 (dehydrogenase/reductase SDR family 11) upregulated in Nigeria and Niger (R-S and C-S comparisons), and in R-S for Chad and Cameroon; AGAP008091 (CLIP-domain serine protease, *CLIPE1*), upregulated in all countries in R-S and C-S comparisons; a chitinase, *Cht5-5*, AGAP013260) upregulated in Nigeria, Nigeria and Chad R-S and C-S, with both induced upregulated in R-C in Chad, as heme peroxidase (*HPX15*), upregulated in R-S and C-S comparisons in Nigeria, Chad and Cameroon, as well as malate dehydrogenase (AGAP000184), upregulated in Nigeria, Chad and Cameroon, R-S and C-S.

However, the most overexpressed genes in Nigeria are a cubilin, a histone (H2B), carbonic anhydrase I, and a cuticular protein, *CPR131* (Additional File 2), while for Cameroon its H2B, Serpin 9 inhibitory serine protease inhibitor, galectin 4 and a Protease m1 zinc metalloprotease.

Several genes were significantly downregulated across the four countries, particularly the cytochrome P450s and the ATP-binding cassette transporters. The most consistently down-regulated P450s were *CYP6AK1* (AGAP010961), downregulated in R-S and C-S comparisons in Nigeria, Niger and Cameroon, while down-regulated in all comparisons in Chad; *CYP4H24* (AGAP013490), downregulated in R-S and C-S comparisons for Nigeria and Cameroon, and downregulated in all comparisons in Niger and Chad; *CYP6M4* (AGAP008214), downregulated in R-S and C-S comparisons in Nigeria, Niger and Cameroon, while downregulated in all comparisons in Chad; and finally *CYP6P5* (AGAP002866), downregulated in R-S and C-S comparisons in Niger and Chad, but only in C-S in Nigeria.

#### Analysis of commonly overexpressed metabolic resistance genes across the Sahel

Special attention was given to the known metabolic resistance genes, implicated in insecticide resistance in *Anopheles* and/or other insects, including, CYP450s, GSTs, carboxylesterases, cuticular proteins, chemosensory proteins/SAPs, uridine diphosphoglucuronosyltransferases, etc, in addition to other important genes, e.g., the immune proteins. Analysis of the data from the list of genes significantly, differentially expressed (FDR-adjusted p < 0.05 and log_2_FC = 1/ FC ≥ 2) revealed the major, commonly upregulated metabolic resistance genes across the Sahel. Top 50 genes from these lists (Additional File 3) from each country are displayed as heatmaps of fold changes in Additional Figure S5a, -b, -c and -d). The most common genes linked with resistance and/or other physiologically important phenotypes are tabulated in Table 1, showing fold changes of 55 genes (R-S and C-S comparisons), common to all countries, or three or two countries at least. The most commonly and consistently overexpressed genes across the Sahel (taking account mean expressions) are the chymotrypsins, -1, -2 and -3 (*CHYM1*/AGAP006709, *CHYM3*/AGAP006711 and *CHYM2*/AGAP006710) (Table 1, Additional Figure S5, Additional File 2, Results section), with fold change for *CHYM1* in R-S and C-S comparisons of 31.62 and 54.41 for Nigeria, 12.31 and 41.59 for Niger, and 18.00 and 18.08 for Chad, and 5.69 and 11.77 for Cameroon; for *CHYM3*, 32.45 and 58.43 for Nigeria R-S and C-S, respectively, 7.30 and 43.27 for Niger, 13.40 and 7.24 for Chad, and 2.59 and 8.15 for Cameroon. Of *Anopheles* metabolic resistance genes, *GSTe2* (AGAP009194) was the most consistently upregulated gene across the Sahel, FC = 9.61 and 7.16, respectively for Nigeria R-S and C-S comparisons; 6.55 and 11.32 for Niger; 9.09 and 10.02 for Chad, 9.68 and 20.67 for Cameroon. Other common GSTs were *GSTZ1*, upregulated in three countries (Nigeria, Niger and Chad), *GSTe4*, *GSTU1*, *GSTD1-4* and *GSTD3*.

**Table 1:** The common detoxification and metabolic genes differentially upregulated in Sahel *An. coluzzii (*log_2_FC values and FDR-adjusted p < 0.05).

| Gene | NIGERIA |  | NIGER |  | CHAD |  | CAMEROON |  | Gene Description |
| --- | --- | --- | --- | --- | --- | --- | --- | --- | --- |
|  | R-S | C-S | R-S | C-S | R-S | C-S | R-S | C-S |  |
| AGAP006400 | 2.2 | 4.2 | 1.7 | 4.5 | - | - | 2.1 | 2.8 | alkaline phosphatase 2 |
| AGAP028491 | 4.3 | 2.8 | 1.4 | 2.7 | 2.9 | 2.6 | - | - | aquaporin, AQP2 |
| AGAP010326 | 14.8 | 11.0 | 12.1 | 21.9 | 14.9 | 1.2 | 14.2 | 17.3 | Aquaporin, AQP3 |
| AGAP005372 | 9.5 | 12.2 | 3.3 | 4.3 | 6.6 | 3.6 | 1.4 | 2.6 | carboxylesterase, COEBE3C |
| AGAP005837 | 4.7 | 5.0 | 1.4 | 3.3 | 1.9 | 2.6 | - | - | carboxylesterase, COEJHE5E |
| AGAP004904 | 3.9 | 2.2 | - | - | 2.3 | 1.7 | - | - | catalase, CAT1 |
| AGAP029127 | 4.1 | 3.2 | - | - | 1.7 | 2.2 | - | - | chemosensory protein 5, CSP5 |
| AGAP006191 | 21.1 | 38.6 | 40.1 | 67.2 | 91.4 | 5.7 | 44.3 | 40.5 | chitinase, Cht24 |
| AGAP013260 | 4.4 | 11.5 | 1.3 | 3.9 | 3.1 | 0.8 | - | - | chitinase, Cht5-5 |
| AGAP006898 | 8.9 | 5.5 | - | - | 2.2 | 3.9 | - | - | chitinase, Cht6 |
| AGAP006709 | 31.6 | 54.4 | 12.3 | 41.6 | 18.0 | 18.1 | 5.7 | 11.8 | chymotrypsin-1, CHYM1 |
| AGAP006710 | 9.9 | 19.4 | 2.1 | 8.8 | 4.2 | 3.9 | - | - | chymotrypsin-2, CHYM2 |
| AGAP006711 | 32.5 | 58.4 | 7.3 | 43.3 | 13.4 | 7.2 | 2.6 | 8.2 | chymotrypsin-3, CHYM3 |
| AGAP000987 | 4.3 | 2.7 | - | - | 1.3 | 2.6 | - | - | cuticular protein, CPAP3-A1b |
| AGAP000988 | 5.3 | 1.6 | - | - | 0.7 | 5.4 | - | - | cuticular protein, CPAP3-A1c |
| AGAP000986 | 7.3 | 4.6 | - | - | 1.7 | 3.5 | - | - | cuticular protein, CPAP3-D |
| AGAP006149 | 3.8 | 0.8 | - | - | 0.7 | 5.8 | - | - | cuticular protein, CPLCX3 |
| AGAP010123 | 33.3 | 54.9 | 2.4 | 13.2 | - | - | - | - | cuticular protein, CPR131 |
| AGAP005456 | 5.2 | 1.8 | - | - | 1.2 | 2.2 | 1.1 | 2.1 | cuticular protein, CPR15 |
| AGAP006009 | - | - | 2.9 | 1.3 | - | - | 3.9 | 1.6 | cuticular protein, CPR30 |
| AGAP009871 | 8.7 | 2.7 | 1.1 | 2.6 | 2.2 | 4.2 | 1.7 | 2.6 | cuticular protein, CPR75 |
| AGAP009874 | 10.6 | 2.2 | 1.1 | 2.6 | 3.7 | 5.6 | 1.1 | 2.0 | cuticular protein, CPR76 |
| AGAP009879 | 7.4 | 2.6 | - | - | 1.9 | 4.6 | - | - | cuticular protein, CPR81 |
| AGAP006422 | 7.5 | 9.4 | - | - | 8.4 | 7.3 | 2.1 | 2.0 | cyanogenic beta-glucosidase |
| AGAP002417 | 5.5 | 4.2 | 0.9 | 1.2 | 1.9 | 1.9 | - | - | cytochrome P450, CYP4AR1 |
| AGAP009246 | 7.4 | 4.6 | 4.0 | 5.6 | 6.0 | 2.6 | 2.0 | 2.6 | cytochrome P450, CYP4C27 |
| AGAP012957 | 5.0 | 2.8 | - | - | 2.9 | 2.5 | 2.0 | 2.6 | cytochrome P450, CYP4D17 |
| AGAP001076 | 5.8 | 4.7 | 1.5 | 1.8 | 3.8 | 2.4 | 0.9 | 1.5 | cytochrome P450, CYP4G16 |
| AGAP000877 | 3.8 | 2.0 | 2.3 | 1.9 | 3.8 | 2.1 | 1.2 | 2.1 | cytochrome P450, CYP4G17 |
| AGAP008358 | - | - | 2.9 | 4.1 | 3.4 | 3.3 | 3.6 | 2.0 | cytochrome P450, CYP4H17 |
| AGAP007480 | 5.4 | 3.9 | 1.3 | 2.7 | 2.5 | 2.0 | - | - | cytochrome P450, CYP6AH1 |
| AGAP002865 | 1.11 | 0.5 | 2.3 | 1.9 | 0.7 | 12.8 | 2.1 | 2.3 | cytochrome P450, CYP6P3 |
| AGAP008207 | 3.6 | 1.9 | - | - | 2.1 | 1.6 | 1.7 | 2.4 | cytochrome P450, CYP6Y2 |
| AGAP008218 | 6.6 | 2.1 | 6.7 | 2.3 | 14.0 | 1.5 | 11.8 | 8.4 | cytochrome P450, CYP6Z2 |
| AGAP008217 | 4.7 | 1.8 | 6.1 | 1.4 | 3.5 | 1.1 | 7.1 | 3.8 | cytochrome P450, CYP6Z3 |
| AGAP000818 | 3.0 | 1.2 | 0.8 | 0.5 | 0.9 | 1.6 | 3.4 | 2.2 | cytochrome P450, CYP9K1 |
| AGAP010400 | 3.6 | 2.6 | 1.6 | 2.8 | 3.3 | 1.4 | - | - | flavin-containing monooxygenase |
| AGAP007920 | 5.7 | 2.4 | 2.8 | 1.8 | - | - | - | - | glucuronosyltransferases, UGT-B19 |
| AGAP004164 | 2.2 | 1.9 | 1.7 | 2.3 | 1.4 | 1.3 | - | - | glutathione S-transferase, GSTD1-4 |
| AGAP009194 | 9.6 | 7.2 | 6.6 | 11.3 | 9.1 | 10.0 | 9.7 | 20.7 | glutathione S-transferase, GSTe2 |
| AGAP009193 | 3.3 | 2.3 | 1.1 | 2.1 | 1.5 | 1.6 | - | - | glutathione S-transferase, GSTe4 |
| AGAP000947 | 4.4 | 2.3 | 1.2 | 2.2 | 1.9 | 3.2 | - | - | glutathione S-transferase, GSTU1 |
| AGAP002898 | 5.5 | 3.5 | 1.7 | 2.6 | 2.8 | 2.4 | - | - | glutathione S-transferase, GSTZ1 |
| AGAP002198 | - | - | 2.0 | 5.8 | 4.8 | 6.6 | - | - | glycine-N-methyltransferase |
| AGAP013327 | - | - | 1.2 | 7.3 | 7.9 | 20.1 | 5.1 | 14.8 | heme peroxidase, HXP15 |
| AGAP002353 | 26.7 | 20.8 | 16.5 | 29.8 | 15.0 | 10 | 14.6 | 14.5 | lipase |
| AGAP000184 | 5.2 | 6.3 | - | - | 4.9 | 4.3 | 1.2 | 1.9 | malate dehydrogenase |
| AGAP008051 | 3.7 | 4.0 | 1.1 | 1.3 | 1.7 | 1.7 | - | - | sensory appendage protein 1, SAP1 |
| AGAP008052 | 5.5 | 10.8 | 1.0 | 1.5 | 3.6 | 1.1 | - | - | sensory appendage protein 2, SAP2 |
| AGAP010815 | 5.7 | 14.4 | 4.1 | 8.6 | 7.8 | 3.8 | 1.7 | 3.1 | thioester-containing protein, TEP1 |
| AGAP008368 | 3.3 | 6.1 | 1.4 | 3.9 | 3.4 | 2.0 | - | - | thioester-containing protein, TEP14 |
| AGAP008654 | 4.6 | 17.4 | 2.1 | 10.0 | - | - | - | - | thioester-containing protein, TEP12 |
| AGAP008227 | 4.1 | 4.0 | 4.0 | 2.0 | 4.6 | 1.5 | 2.6 | 4.1 | trehalose 6-phosphate synthase/phosphatase |
| AGAP012818 | 22.1 | 14.6 | 31.1 | 25.4 | 6.0 | 4.3 | 44.9 | 43.4 | V-type proton ATPase, subunit a |

Other most consistently and highly upregulated genes include a lipase, AGAP002353: FC of 26.73 and 20.80 for Nigeria R-S and C-S comparisons, 16.53 and 29.76 for Niger, 14.95 and 9.95 for Chad, 14.58 and 14.53 for Cameroon; a chitinase, *Cht24* (AGAP006191), FC = 21.12 and 38.55 for Nigeria R-S and C-S, 40.08 and 67.18 for Niger, 91.42 and 5.71 for Chad, 44.26 and 40.53 for Cameroon); a V-type protein ATPase subunit a (AGAP012818, FC = 22.12 and 14.61 for Nigeria, 31.05 and 25.43 for Nigeri, 6.00 and 4.31 for Chad, 44.92 and 43.4 for Cameroon), and an aquaporin, *AQP3* (AGAP010326, FC = 14.81 and 11.03 for Nigeria, 12.09 and 21.94 for Niger, 14.86 and 1.21 for Chad, 14.17 and 17.3 for Cameroon).

The most overrepresented gene family, and most consistently upregulated across the Sahel were the cytochrome P450s, with *CYP6Z2*, *CYP6Z3*, and *CYP4C27*, in addition to the two P450s linked to cuticular resistance, *CYP4G16* (upregulated in Nigeria and Chad) and *CYP4G17* (upregulated in Nigeria, Niger and Chad). The three P450s, *CYP6Z2, CYP6Z3 and CYP4C27* are consistently overexpressed: FC for *CYP6Z2* is 6.55 and 2.10 for Nigeria R-S and C-S comparisons, 6.73 and 2.32 for Niger, 13.96 and 1.48 for Chad, 11.84 and 8.39 for Cameroon; FC for *CYP6Z3* is 4.73 and 1.82 for Nigeria, 6.09 and 1.42 for Niger, 3.51 and 1.11 for Chad, 7.11 and 3.79 for Cameroon; and FC for *CYP4C27* is 7.36 and 4.6 for Nigeria, 4.03 and 5.64 for Niger, 6.04 and 2.62 for Chad, 2.01 and 2.63 for Cameroon. The two well-known insecticide resistance genes *CYP6P3* and *CYP6M2* were not overexpressedprominently across the Sahel of these countries.

The second most commonly overrepresented gene families belong to cuticular proteins, including cuticular proteins Rebers and Riddiford (RR), several cuticular proteins of low complexity (CPLC) and chitin-binding cuticular proteins (CPAP), with the commonly upregulated ones across all four countries being *CPR75* (AGAP009871) and *CPR76* (AGAP009874).

Two carboxylesterases picture prominently across the Sahel – the beta esterase, *COEBE3C* (AGAP005372) upregulated in all four countries, while *COEJHE5E* was common to Nigeria, Niger and Chad. For Phase II metabolism enzymes, several uridine-diphospho-glucuronosyltransferases, UGTs were upregulated, including the *UGT-B19*, which is upregulated in Nigeria and Niger (Table 1), and unknown UGT (AGAP006222) upregulated across all the countries (Additional File 2).

Other genes in the top 55 metabolic genes include the malate dehydrogenase, *TPS* (*TPS1/2*), *AQP2*, thioester-containing proteins, *TEP1, -12* and *-14,* a cyanogenic-beta-glucosidase (AGAP006422) and three chemosensory proteins, *CSP5*, sensory appendage proteins, -1 and -2.

### Gene Ontology Enrichment Analysis

Gene ontology enrichment analysis, for genes significant in comparisons (log_2_FC = 1, p < 0.05) revealed crucial differences in overrepresented GO terms between the up- and - downregulated genes, in data from the four countries. For example, the most over-represented, semantically similar GO terms associated with xenobiotics metabolism is oxidoreductase activity ((highest frequency of 12.9 %, Additional Figure S6a, REVIGO Table View), which cluster together in Nigerian R vs S comparison (for genes upregulated in R). Other over-represented GO terms in this comparison include glutathione S-transferase activity (the most enriched/specific term), peroxidase activity, odorant binding and chitin binding. In contrast, GO terms over-represented in R vs S (down-regulated in R) were mostly involved in neurotransmission, metal ions binding and receptor channelling activities (Additional Figure S6b).

For R vs C comparison (upregulated in C) the over-represented GO terms include oxidoreductase (frequency = 12.88 %), glutathione S-transferase activity (the most enriched/specific term), peroxidase activity, glucuronosyltransferase activity, aldehyde oxidase activity, chitin binding, carbohydrate binding activities, etc, (Additional Figure S6c). In contrast the most enriched GO terms, downregulated in C were endopeptidase activities, Toll binding, oxidoreductase (frequency = 1.21 %) and monooxygenase activities (Additional Figure S6d). For the rest of the three countries similar contrasts were also observed, between phenotypes, and detailed in Additional File 1, Results section.

### Quantitative PCR validation of expression profiles of metabolic resistance genes

The relative expression levels of 12 metabolic resistance genes were validated. The qRT-PCR results support the transcriptomic patterns obtained from the RNAseq analysis, with *GSTe2* as the most overexpressed gene, followed by *CYP6Z2* (Figure 3). For example, for R-S and C-S comparisons *GSTe2* had fold changes of 49.25 ± 5.05 and 28.45 ± 4.45, for Nigeria (*p* ≤ 0.0001); 37.01 ± 2.22 and 33.34 ± 6.61 for Niger (*p* ≤ 0.0001); 29.97 ± 5.39 and 30.16 ± 6.16 for Chad (*p* ≤ 0.0001); and 28.30 ± 3.30 (*p* ≤ 0.0001) and 17.03 ± 7.03 for Cameroon (*p* ≤ 0.001). *CYP6Z2* has R-S and C-S fold changes of 15.93 ± 2.54 (*p* ≤ 0.001) and 7.35 ± 1.52 (*p* ≤ 0.05), for Nigeria; 24.21 ± 2.17 (*p* ≤ 0.001) and 20.83 ± 3.02 for Niger (*p* ≤ 0.001); 15.02 (*p* ≤ 0.001) ± 2.01 and 6.09 ± 2.08 for Chad (*p* ≤ 0.05); and 10.58 ± 3.00 (*p* ≤ 0.01) and 3.43 ± 1.43 for Cameroon. Correlation analyses for R-S comparisons for all genes revealed positive and significant association (Additional Figure S7) between the RNA-seq and qRT-PCR data in data from Niger (R^2^ = 0.58, p = 0.03) and Chad (R^2^ = 0.408, p = 0.025), with positive, but non-significance seen in Cameron (R^2^ = 0.58 p = 0.08) and Nigerian (R^2^ = 0.503, p = 0.06).

**Figure 3:** Validation of candidate resistance genes. qRT-PCR profile of twelve metabolic resistance genes (in R-S and C-S comparisons) from the Sahelian *An. coluzzii*. Each bar (data point) is analysis from three biological replicates and three technical replicates for each gene, with error bars representing standard deviations. Significant differences are also indicated with the symbols ϯ, Ϯ, † and ‡, for p < 0.0001, p < 0.001, p < 0.01 and p < 0.05, respectively.

### Detection of Signatures of Selective Sweeps

Signatures of selection were investigated in the major metabolic resistance genes, by estimating Tajima’s D per gene within populations, and *F*st per gene between population pairs. These tests of neutrality revealed several genes exhibiting genetic differentiation, or possibly undergoing expansion. Among the top 100 most overexpressed metabolic genes across the four countries (Additional File 2), 6 genes were possibly undergoing genetic differentiation. These include *TEP1,* and *TEP3*, with average Tajima’s D of -1.4 and -1.6 respectively, in populations from Chad, and *F*st values of ≥ 0.85 for *TEP1*, for Nigeria, Chad and Cameroon samples, compared to the average *F*st of 3L chromosomal, which was calculated as 0.24 (Additional File 3), *CYP9K1* (average Tajima’s D = -1.00 for Cameroon, and *F*st within ranges of 0.86 - 1 for Nigeria, Cameroon and Niger, compared with 0.21 calculated for the chromosome X), *CPR15* (average Tajima’s D = -0.94); *CPAP3-A1b* (Tajima’s D = -0.70 in Chad, -0.95 in Nigeria, and -1.32 in Cameroon, with *F*st values in range of 0.7 - 0.9 for the four countries, compared with the average *F*st of 0.21 for chromosome X) and *GSTU1*. Several genes from the GST family exhibited strong genetic differentiation, with reduced variations prominently in *GSTe2* (< 3 SNPs in Chad and Cameroon samples), with Tajima’s D of -1.45 and -1.15 for Nigeria and Niger, and combined. Information on more genes is provided in Additional File 1, and detailed information on all genes can be found in the Additional File 3.

### Identificaton of genetic variants associated with insecticide resistance

The RNA-Seq-Pop workflow calculated allele frequencies of variants of interest found in the raw RNA-sequencing read data. Figure 4 displays allele frequencies of each variant in respective treatment replicates used for the RNAseq, as well as Ngoussou. Several mutaions were found within the voltage-gated sodium channel. In addition to the L995F *kdr* mutation, the recently identified V402L and I1527T mutations exist in all the field populations across the Sahel, at frequencies ranging from 29 % to 67 %. We also detected the N1570Y mutation, which shares the same haplotype as L995F. The G280S *acetylcholinesterase*-1 mutation was also observed in low to moderate frequencies in all populations except for Ngoussou and samples from Niger. The L43F pyrethroid resistance marker of *CYP4J5* was also seen, fixed in most populations.

**Figure 4:** Identification of resistance variants of interest. A heatmap showing frequencies of the resistance variants (haplotypes) in some key genes of interest.

### Detection of chromosomal inversion polymorphisms and metabolic genes within its breakpoints

The frequency of the major inversion polymorphisms in chromosome 2 were calculated, considering ploidy (8 individual female mosquitoes were pooled for RNA extraction, for each replicate). Additional Table 5 provided the frequencies of the respective inversions for the populations from each country, as well as for Ngoussou. High frequency of the 2La, 2Rb and 2Rc were observed, in contrast to the 2Rd, 2Rj and 2Ru which were in lower frequencies [except for the 2Rj in Nigeria (30.38 %) and 2Ru in Cameroon (24.29 %)]. The 2La inversion was found fixed in the field populations (100 % in Nigeria and Cameroon, 99.92 % in Niger and 99.21 % in Chad), in contrast with Ngoussou, with frequency of only 6.26 %. Similar pattern was observed with the 2Rb inversion, with high frequencies in three field populations (79.01 %, 84.85 %, 80.22 % in Nigeria, Niger and Chad), but lower in Cameroon (29.08 %) and Ngoussou (5.35 %). Frequencies of the 2Rc inversion were 88.54 %, 85.52 %, 88.82 %, 46.99 % and 9.89 % for Nigeria, Niger, Chad, Cameroon and Ngoussou respectively.

Some genes among the top 100 most overexpressed metabolic resistance genes (above) and which were likely undergoing recent population expansion were located within the 2La, 2Rb and 2Rc inversions (Additional File 4). These include the *hsp83* (AGAP006958) and molecular chaperone hptG (AGAP006961), *CPLCX3* (AGAP006149), *CPLCA1* (AGAP006145,) within the inverted 2La chromosomal arm. Other genes within 2La inversion include a group of chitinase enzymes (*Cht6*, *-8*, *-11* and *-24*), most notably the *Cht24* and *Cht6*, which are among the top 50 most overexpressed genes in all countries. Other genes include: *COEJHE5E*, one of the 14 carboxylesterases in the 2La region, and which was among the top 100 most overexpressed genes across the Sahel; the most overexpressed chymotrypsin genes, *CHYM1* and *CHYM2*; three ionotropic receptors, *IR136* (AGAP006440), *IR139* (AGAP006691) and *IR142* (AGAP006407); as well as five P450s, *CYP301A1*, *CYP302A1*, *CYP4J5*, *CYP4J9* and *CYP4J10*. The most overrepresented genes within 2La inversions are the CPR cuticular proteins (73 in total), with *CPR21*, *-26*, *-30*, *-59* and -*140* from the list of the top 100 resistance genes from the Sahel.

Carbonic anhydrase I (the most overexpressed metabolic resistance in Nigeria, AGAP013402) sits within 2Rb inversion. Four chitinases (*Cht4*, *Cht5-1*, *Cht5-3,* and *Cht5-5*) are located within 2Rb inversion; *Cht5-1* and *Cht5-5* were among the observed top 50 most overexpressed metabolic genes in the Sahel. Lipase (*AGAP002353*) which is the among the top 6 overexpressed metabolic gene in the four countries sit within the 2Rb inversion. Eight CYP450s reside in the 2Rb inversion, including *CYP4D15*, *CYP4D17* and *CYP4AR1*, which were among the top 100 metabolic resistance genes. Three cuticular proteins, *CPR7*, *-8* and *-9* were also within the 2Rb inversion.

Interesting genes sitting within 2Rc inversion include *GSTZ1*; 5 ionotropic receptor genes, *IR7i* (AGAP013363), *IR7u* (AGAP013285), *IR7t* (AGAP002763), *IR7w* (AGAP013416) and *IR41a* (AGAP002904); a carboxylesterase, *COEAE6O* (AGAP002863); and 11 CYP450s, including *CYP6AA1* (AGAP002862), *CYP6AA2* (AGAP013128), *CYP6P15P* (AGAP002864), *CYP6P3* (AGAP002865), *CYP6P5* (AGAP002866), *CYP6P4* (AGAP002867), *CYP6P1* (AGAP002868), *CYP6P2* (AGAP002869), *CYP6AD1* (AGAP002870) and *CYP6Z4* (AGAP002894)

### Investigation of the polymorphism in the coding sequences of *GSTe2* and *CYP6Z2*

Analysis of the polymorphism patterns of full-length cDNA sequences of *GSTe2* (666 bp) and *CYP6Z2* (1479 bp) from the Sahel region of Africa revealed complete homogeneity for *GSTe2*, with no polymorphism detected in the field populations (all sequences were identical to those from Ngoussou and the AGAP009194 reference). This suggests a fixed allele, consistent with the observation from the analyses from the fixation index (Additional File 3). For *CYP6Z2* (1479 bp), homogeneity was observed within each country and Ngoussou, except for Niger, characterised by an unusually high polymorphism (Additional Figure S8a, -b). *CYP6Z2* is polymorphic with 10 haplotypes across the Sahel, with 75 polymorphic sites of which 65 were synonymous, and 12 led to amino acids substitutions. The bulk of the polymorphisms were contributed from larger variations in the Niger and Cameroon (S = 42 and 23 respectively), while highest homogeneity was observed in Chad, with a single haplotype. Haplotype diversity is high (H_d_ = 0.921), from 10 haplotypes out of only 20 sequences, with the lowest H_d_ in the Chad sequences, suggesting a directional selection/fixed allele. The haplotypes cluster according to origin on the maximum likelihood phylogenetic tree, except for Niger (Additional Figure S8c).

### Investigation of the role of intergenic region elements in overexpression of *GSTe2* **and *CYP6Z2***

#### Investigation of polymorphism in the intergenic region/regulatory elements

To investigate polymorphisms in the regulatory elements of the above genes, the 351 intergenic regions of *GSTe2* (spanning the 43 bp 3’-UTR of *GSTe1,* 248 bp flanking sequence and 60 bp 5’-UTR of *GSTe2*) preceding the start codon were amplified from 10 each of DDT-alive and -dead females from the 4 Sahel countries, as well as from Ngoussou females, successfully. For *CYP6Z2,* 1078 bp (spanning 38 bp 3’ UTR of *CYP6Z1,* 937 bp flanking sequence and 103 bp 5’UTR of *CYP6Z2,* preceding the start codon of *CYP6Z2*) were used to amplify fragments from 10 each of deltamethrin-alive and -dead females from Nigeria, and from Ngoussou females.

Out of the 90 *GSTe2* 5’-UTR sequences analysed differences were observed between the alive and dead mosquitoes, with a total of 65 sequences from alive and dead females (regardless of country of origin) identical to the 10 sequences of Ngoussou. From the 40 sequences of the dead females, 39 were identical to Ngoussou (regardless of the country). Twelve sequences, all from the alive females (from across the four countries) were similar, with several mutations in putative transcriptional factors binding sites, which may impact overexpression of the *GSTe2.* In short, 8 mutations were shared in common between these 12 sequences of DDT-alive females from across the countries (3 each from Nigeria, Chad and Cameroon) and additional mutations in the Niger samples (4 sequences). These mutations include (i) T->A transition withing the cellular myb-DNA (c-myb) binding domain (Additional Figure S9), (ii) a T->C transversion in the zinc-finger homeodomain, *δ*EF1 (*δ* elongation factor 1) binding site, (iii) T->A transition in the nuclear matric protein 4 (NMP4), (iv) simultaneous insertion of adenine and a transition T -> C, in positions 113 and 114 respectively, between the Fork-head box L1 and c-EST binding sites, (v) an A -> C transversion in a second NMP4, (vi) a T -> C transition, six nucleotides downstream the nuclear factor *κ*B (NF-*κ*B), (vii) followed by an A -> G transversion 3 nucleotides downstream the NF-*κ*B, (viii) a G -> A transversion, 7 nucleotides upstream the GC box, (ix) a C -> G transition, 2 nucleotides downstream the arthropod initiator (Inr consensus) sequence, and finally (x) a C -> G transition, within the 5’-UTR of the *GSTe2*, 28 nucleotides downstream the transcriptional start site (49 nucleotides upstream the *GSTe2* start codon).

Analysis of the 90 sequences revealed a very low polymorphism in the dead mosquitoes (S = 0, for Niger-dead, Chad-dead and Ngoussou), but high polymorphism in the alive (S = 16 for Nigeria-alive, and 13 each for Chad-alive and Cameroon-alive) (Additional Table 6). All sequences produced 17 polymorphic sites, with 6 haplotypes (Additional Figure S10a and - b). High haplotype diversities were obtained from the alive mosquitoes (for example, Nigeria alive, Hd = 0.71, Niger-, Hd = 0.53). Regardless of country of origin, the haplotypes cluster according to phenotype on the maximum likelihood phylogenetic tree, with the alive haplotypes forming a distinct (separate) clade (Additional Figure S10c).

With regard to *CYP6Z2* no major differences were observed when comparing the deltamethrin-alive and -dead sequences with the Ngoussou.

#### Measurement of activities of the 5’ regulatory elements of the GSTe2 and CYP6Z2

Initial promoter analyses were conducted with the 5’-regulatory element sequences of *GSTe2*, for the DDT-alive females (representative sequence with the 8 common polymorphic positions, designated, Sahel-alive), comparing it with the sequence from the DDT-dead/Ngoussou (designated Ngoussou/Sahel-dead). For *CYP6Z2*, a predominant sequence from the Nigeria field sample (with 8 nucleotide insertion) was compared with the Ngoussou sequence.

The ability to drive heterologous expression of the firefly luciferase was determined, with the Sahel-alive construct and Ngoussou/Sahel-dead producing increased luciferase activity [∼3090× (normalized luciferase activity = 2.58) and 479× (normalized activity = 0.400), respectively] compared with the promoterless pGL3-Basic vector. But the Sahel-alive construct was significantly more active than the Ngoussou/Sahel dead counterpart [promoter activity ∼6-fold higher (Tukey HSD Q = 8.11, p = 0.004)]. In contrast, for *CYP6Z2* no significant differences were observed when comparing the field construct (normalized luciferase activity = 5.95 for alive/dead field construct), compared with 5.84 for the Ngoussou construct.

#### GSTe2 promoter delineation and measurement of activity

Sequential deletion of the intergenic region of the *GSTe2* resulted in progressive reduction in luciferase activity. Deletion of the 43 bp 3’-UTR of the *GSTe1* (-308 from the start codon of *GSTe2*) reduced activity of the Sahel-alive construct by only 18.6 % suggesting that the c-Myb transcriptional factor binding site may not be critical for overexpression (Figure 5). Deletion of an additional 38 nucleotides from the flanking region (-270 fragment, obliterating the δEF1 binding site) had comparable impact as above, with activities reducing by 15.3 % only. But removing the *GSTe-1* 3’-UTR and an additional 46 nucleotides from the flanking region (-262 fragment, which obliterated in addition the FOX-L1 putative binding site) reduced activity by 64.1 % (Q = 6.18, p = 0.01) indicating the importance of this binding site. Deletion of *GSTe1* 3’-UTR plus 75 nucleotides of the flanking region (-232 fragment, located 10 nucleotides from the c-EST binding site) significantly reduced the activity by ∼70% (Q = 6.14, p = 0.01), suggesting the importance of the simultaneous insertion of adenine and transition T -> C, in positions 113 and 114 respectively, between the FOX-L1 and c-EST binding sites. Removing the fragment of the 5’UTR of *GSTe2* (43 nucleotides preceding the AUG codon obliterated the promoter activity, reducing the luciferase expression by 98 % (Q = 9.41, p = 0.002). This is despite presence of all the above binding sites and transcriptional start site in this fragment. Deletion of the 5’-UTR from Ngoussou significantly reduced activities as well (reduction by 88 % compared with the full Ngoussou intergenic region construct, Q = 13.75, p = 0.001). Overall, these findings suggest that the essential binding sites for overexpression of *GSTe2* span the FOX-L1 and c-EST binding sites, with the 5’-UTR essential for activity.

**Figure 5:** Characterization of intergenic region (5’ regulatory element) of *GSTe2*. Results of dual-luciferase reporter assays of the promoter (intergenic region) constructs, showing progressive loss of activity following sequential deletion of the constructs.

### Investigating the role of *GSTe2* and *CYP6Z2* in insecticides resistance using transgenic analysis

A qRT-PCR was conducted using the transgenic flies to first establish overexpression of the above genes. Relative fold changes (FC) of 32.7 ± 4.6 and 25.30 ± 2.8 were obtained in flies overexpressing the *GSTe2* and *CYP6Z2* respectively, compared with control flies (progenies of crosses between the parental line flies with no gene insertion, crossed with GAL4/UAS driver line) (Additional Figure S11).

Contact bioassays carried out using 0.05 % α-cypermethrin revealed a high susceptibility in the transgenic flies expressing *GSTe2* (Act5C-GAL4-UAS-GSTe2) and controls (Figure 6a), with mortalities increasing from 62 % and 70 % respectively in 1 h, to 94 % and 99 % in 24 h. However, significantly reduced mortalities were observed in transgenic flies expressing *CYP6Z2* (Act5C-GAL4-UAS-CYP6Z2) compared to control flies, at 1 h (mortality = 35 % *vs* 70 % in control, p < 0.001) and 3 h (mortality = 52 % vs 89 % in the control, p < 0.001). High susceptibilities were also seen in all the experimental flies exposed to 0.05 % deltamethrin (both for *GSTe2* and *CYP6Z2* flies), except for 1 h with flies expressing *CYP6Z2* (mortality = 47 % compared with 58 % in the control flies, p < 0.010).

**Figure 6:** Validation of the role of metabolic resistance genes in insecticide resistance. Results of insecticide susceptibility bioassays with transgenic flies expressing *GSTe2* and *CYP6Z2*. **a.** α-cypermethrin and deltamethrin, **b.** permethrin, **c.** 3-phenoxybenzaldehyde and 3-phenoxybenzyl alcohol, **d.** DDT and clothianidin.

The *GSTe2* transgenic flies were highly resistant to 2 % permethrin, with no mortality at all in 1 h (Figure 6b), and average mortalities of only 8.5 % at 24 h (p < 0.0001), compared with 85 % at 1 h for control flies, which increased to 100 % from 3 h.

Initial exposure to 4 % of 3-phenoxybenzaldehyde (PBAld) and 3-phenoxybenzyl alcohol (PBAlc) had no toxic effect on *CYP6Z2* transgenic flies (Figure not shown). However, 5x concentration of these primary products of pyrethroid hydrolysis induced mortalities, albeit low in all flies (Figure 6c). For PBAld, mortalities ranged from 15.5 % and 17.5 % at 1 h, for *CYP6Z2* transgenic flies and control flies respectively, to 30 % and 27.5 % at 24 h. Although low mortalities were observed with PBAlc, but at 1 h and 3 h exposure times the mortalities in the transgenic flies expressing *CYP6Z2* were significantly lower compared with mortalities from the control flies (1 h mortality = 2.5 % *vs* 12.5 %, p < 0.001; 3 h mortality = 9.5 % *vs* 17.5 %, p < 0.001).

Marginal tolerance towards DDT was observed in the *CYP6Z2* transgenic flies when compared with the control flies, but only at 3 h (p < 0.001) and 6 h (p < 0.01) (Figure 6d). This is in contrast with the *GSTe2* transgenic flies, with very low mortalities, in ranges of 0 % to 9.52 % for 1 h to 24 h, when compared with control flies (mortality = 34 % in 1 h and 100 % in 6 – 24 h, p < 0.0001).

Susceptibility to clothianidin was very high (Figure 6d), but surprisingly the *CYP6Z2* transgenic flies exhibited a contrasting phenotype, with significantly higher mortalities compared with the control flies, at 1 h (mortality = 67 % vs 42 %, p < 0.001), 3 h (mortality = 71 % vs 53 %, p < 0.001) and 6 h (mortality = 78 % vs 62 %, p < 0.01).

Taken together, these results confirmed that over-expression of *GSTe2* alone is sufficient to confer resistance to type I pyrethroid (permethrin) and DDT, while overexpression of *CYP6Z2* alone, may confer marginal resistance to the α-cypermethrin and PBAlc.

## Discussion

Escalating insecticide resistance across Africa (7–9, 52), if not tackled will compromise the malaria control and elimination efforts. Molecular markers of metabolic resistance e.g., (10, 11, 21), will support evidence-based control and resistance management. Identification and validation of resistance markers in malaria vectors across regions of sub-Saharan Africa can promote communication, cooperation and coordination among malaria control/elimination programs, and allow control efforts to be tailored to the vector species involved in transmission across borders (53), and tracking of the evolution and spread of resistance markers across regions (22). To support malaria pre-elimination efforts in Africa, in this study we targeted the Sudan savannah and Sahel (regions of Africa sharing similar eco-climatic conditions, and characterised by high seasonal transmission), which are ideal for control and elimination of malaria using seasonal vector control (25) and chemoprevention.

### *Anopheles coluzzii* is a major malaria vector across the Sudano-Sahelian transects

Contrary to the previous observations that *An. arabiensis* tends to predominate in arid savannas, while *An. gambiae* is the dominant species in humid forest zones (54–56), *An. coluzzii* has repeatedly been identified as the major malaria vector in the Sudan savannah and Sahel of several, neighbouring countries (including in northern Nigeria, southern Niger, central Chad, and northern Cameroon) (7, 26–28, 57) in recent years, suggesting that this vector has adapted well in drier regions of the Sahel transects, and is probably predominating over *An. arabiensis* and *An. gambiae s.s*. This is not surprising as this species exhibits higher exploitation of breeding sites associated with anthropogenic activities, and behavioural plasticity to avoid predators (58), and is known to survive a long dry season *in situ*/aestivation, which allows it to predominate, becoming the primary force of malaria transmission (59, 60).

### Common metabolic genes mediate multiple resistance in the Sahelian *An. coluzzii*

Several genes shown to confer metabolic resistance in *Anopheles* mosquitoes and other insects were found constitutively overexpressed and/or induced in this study. *GSTe2* (AGAP009194) is one of the most regularly encountered metabolic genes in resistant populations of the major malaria vectors *An. gambiae*, *An. coluzzii* and *An. funestus* (20–23). *Anopheles gambiae GSTe2* has been validated, using transgenic flies to confer DDT (20) and fenitrothion (61) resistance. It was extensively studied in *An. funestus*, in which it was shown to confer cross-resistance to DDT and permethrin (22), reduce efficacy of the LLINs, PermaNet 2.0 and PermaNet 3.0 (side panels) (23), and even increase the longevity of the resistant populations carrying the 119F mutation (62). These and our findings suggest that the overexpression of this GST alone can confer resistance to three insecticides from three different classes (DDT, permethrin and fenitrothion). The absence of mutations in the cDNA coding sequences of *GSTe2* in *An. coluzzii* from these four countries, suggest that overexpression of this GST is enough to confer resistance.

Several cytochrome P450s previously linked with insecticide resistance were found overexpressed across the Sahel. For example, *An. gambiae CYP6Z2* (AGAP008218) known to metabolize carbaryl (63), the insect juvenile hormone analogue insecticide, pyriproxyfen (64) and mitochondrial complex I inhibitors, fenazaquin, pyridaben and tolfenpyrad (65). This P450 also plays a pivotal role in the clearance of pyrethroid insecticides via further catabolism of pyrethroid derivatives (PBAld and PBAlc) obtained by the action of carboxylesterases (66), in line with our findings of this gene conferring marginal tolerance to high concentration of PBAlc, and α-cypermethrin. However, our findings suggest that overexpression of this P450 may enhance the efficacy of clothianidin, which will be epidemiologically advantageous in terms of control. Indeed, bioactivation by P450s is known to be a requirement for insecticidal toxicity of several classes of insecticides, e.g., the organophosphates and chlorpenafyr. In contrast to *GSTe2*, the *CYP6Z2* from across Sahel contain three cDNA mutations, which makes it different from Ngoussou coding sequences. These are K^211^N and T^218^S mutations, both within the substrate recognitions site 2, and an A^282^E. Other important CYP450s found to be commonly overexpressed across the Sahel include the *CYP4C27* (AGAP009246), *CYP6Z3 (*AGAP008217) and *CYP9K1* (AGAP000818), all three shown to be consistently overexpressed in field populations of *An. gambiae/coluzzii* and *An. funestus* across Africa (9, 67, 68). *CYP9K1* has been shown to be epidemiologically important pyrethroid-metabolising P450 linked with metabolism of deltamethrin and pyriproxyfen in *An. gambiae* (69). We also have recently shown that his P450 is involved in pyrethroids in *An. funestus* (70).

Several carboxylesterases were also upregulated/induced across the Sahel, with the *COEBE3C* (AGAP005372) upregulated in all four countries. This beta esterase is enriched in the legs (where xenobiotic detoxification probably occur) of pyrethroid-resistant *An. coluzzii* (71). Other genes consistently overexpressed across the Sahel, include the chymotrypsins (*CHYM3*/AGAP006711 and *CHYM1*/AGAP006709) and a lipase (AGAP002353). Chymotrypsins are known to defend insects against plants’ proteinase inhibitors (72); and previous transcriptional studies have shown *CHYM1* and *CHYM3* overexpressed in insecticide resistant populations of *An. gambiae* and *An. coluzzii,* respectively (9, 67). Using *in vitro* and *in vivo* tools, lipases have been linked with deltamethrin resistance in *Culex pipiens* pallens (73).

### Common metabolic resistance markers probably exacerbate resistance across the Sahel

Several well-known genetic variants implicated in resistance, as well as the recently discovered ones exist in high frequencies in the Sahelian *An. coluzzii,* compared with the Ngoussou. For example, the pyrethroid resistance *CYP4J5*-L43F marker (74) was found fixed across the Sahel. The G280S/G119S *ace*-1 mutation, found in high frequencies across Sahel confers organophosphate an carbamates resistance (75), and is shown recently to confer resistance to pirimiphos-methyl, in *An. coluzzii*/*gambiae* (76). Several mutations found within the VGSC have recently been described/validated. For example, the resistance mutation, L995F (77) and the V402L/I15227T haplotypes have been observed across Africa (78). Recently, the two mutations (V402L/I15227T) are described to be in tight linkage and mutually exclusive to the classical L995F/S mutations (77). Our results suggest haplotypes carrying the V402L/I1527T combination plus the L995 replacement do exist in the Sahel *An. coluzzii*. Not only that, it is also in addition to the N1570Y replacement. However, isolation of Ngoussou colony over generations in the artificial insectary conditions could have led to genetic isolation and drift, from inbreeding, leading to strong population differentiation in comparison to the natural populations studied here.

In contrast to *An. funestus GSTe2*, where overexpression and 119F mutation combined to confer extreme DDT resistance (22), the absence of amino acid replacements in the *An. coluzzii GSTe2* from the Sahel suggests that overexpression alone is the key mediator of DDT and permethrin resistance. This is supported by the higher activity in the regulatory regions, harbouring an insertion and nucleotide substitutions in the alive mosquitoes. Indeed, some of the mutations we have found within the intergenic region of *GSTe2* are similar to those observed in a previous study (79).

### Cuticular resistance mechanism probably playing a key role in Sahelian *An. coluzzii*

Our results suggest cuticular mechanism play a role in pyrethroid resistance in these populations. For example, the findings of *CYP4G16* (AGAP001076) and *CYP4G17* (AGAP000877) overexpressed across the Sahel. The former P450 was previously shown to be involved in epicuticular hydrocarbon biosynthesis associated with resistance (80).

The three major classes of the insect cuticular proteins - the CPR, CPLC and CPAP, were found overrepresented in the top overexpressed genes across the Sahel, with the commonly upregulated ones being *CPR76*, *CPR15* and *CPR30*, a chitin-binding cuticular protein, *CPAP3-A1b*, and cuticular proteins of low complexity, *CPLCX3* and *CPLCA1*. Indeed, *CPAP3-A1b* (AGAP000987) have been shown to be highly overexpressed in deltamethrin-resistant Sahelian population of *An. coluzzii* from Burkina Faso (9) and induced by blood feeding in *An. gambiae* (81). The CPR and CPLC cuticular proteins have been described to potentially play a crucial role in insecticide resistance through leg cuticle remodelling/thickening, regulating penetration rate of insecticides in *An. coluzzii* (71). Furthermore, a recent study has found most of the cuticular proteins we have described here, as highly overexpressed in permethrin/malathion resistant populations of Ethiopian *An. arabiensis* (82), e.g., *CPR30*, *CPR75*, *CPR81*, and *CPLCP11*. Out of the several chitinases overexpressed across the Sahel, four were amongst the top 50 most overexpressed metabolic genes. These include the *Cht24* and *Cht6* that have been shown to be overexpressed in the *An. arabiensis* from the above study (82), with the ortholog of *Cht24*, AARA007329 among the top 10 most overexpressed genes in *An. arabiensis*, in line with our observations in *An. coluzzii* across the Sahel. There is an overwhelming need to functionally investigate the role/contribution towards insecticide resistance of these cuticular proteins, chitinases and a chitin synthase (AGAP001748) significantly overexpressed in the field *An. coluzzii* from Nigeria and Chad.

### Insecticide resistance- and thermotolerance-associated genes sit within chromosomal inversions

In this study, the findings of high frequency of 2La, 2Rb and 2Rc inversion polymorphisms in the populations of *An. coluzzii*, compared with the Ngoussou, suggested strong phenotypic adaptations in this species, across the Sahel. Most importantly, in addition to several of the cuticular protein genes associated with resistance (chitinases, chitin synthase, CPR, CPLC and CPAP proteins), several other genes previously implicated in thermotolerance and/or desiccation resistance in *An. gambiae*/*coluzzii,* and which were highly overexpressed in this study sit within these inversions. For example, the heat shock proteins, *hsp83* (AGAP006958) and *hsp90* hptG (AGAP006961), both of which are known to be heat- and insecticide-stress inducible (83) and were among the core set of hsp genes involved in a common and immediate response to thermal stress in *An. gambiae* populations (84), sit within the 2La inversion. These two genes were among the overexpressed genes in both heat-hardened and permethrin-resistant *An. coluzzii* populations from northern Nigeria (57). Several ionotropic glutamate receptors were found within the 2La inversion breakpoints: *IR136* (AGAP006440), *IR139* (AGAP006691) and *IR142* (AGAP006407). This is not surprising as ionotropic receptors are commonly associated with chemosensation, thermosensation, and hygrosensation (85, 86), characteristics which can confer adaptive advantages in xeric environs. The *IR25a* (AGAP010272) and *IR21 a* (AGAP008511) are known to mediate both humidity and temperature preference in the fruit fly, *D. melanogaster* (86, 87), in addition to *IR21a* driving heat seeking and heat-stimulated blood feeding in *An. gambiae* (87). These two genes have been shown to be overexpressed/induced in thermotolerant/permethrin-resistant populations of *An. coluzzii* (57).

## Conclusions

Information on molecular basis of resistance and/or resistance genes and its markers facilitates evidence-based control measures. In this study we characterised a major malaria vector, *An. coluzzii* from the Sahel region of four countries, with findings which could promote evidence-based, cross-border policy towards local and regional malaria control. The study found that across Sahel (where malaria is highly seasonal, reaching its peak in the rainy season), *An. coluzzii* is a dominant vector. And that a handful of common cross-resistance genes are responsible for multiple insecticide resistance in this species. Findings from this study suggest pleotropic role of some key genes – able to confer insecticide resistance and/or stabilize the insecticide resistance gene, at the same time conferring environmental adaptations, such as the ability to survive thermal stress (thermotolerance), as expected in this Sahelian region. From operational vector control perspective this study provided evidence of the role of key insecticide metabolism gene, *CYP6Z2* in increasing insecticidal potency of clothianidin, which could increase the efficacy of the ingredients in malaria control tools, when targeting field populations overexpressing this key P450.

## Supporting information

Supplementary Text - Methods and Results

Common differentially expressed genes across the Sahel

Population genetics analysis

Genes within 2La 2Rb and 2Rc chromosomal inversions

## Declarations

### Ethics approval and consent to participate

This study did not use human participants, human data or human tissue. Ethical approval for collection of indoor resting female mosquitoes in Nigeria, Niger, Chad and Cameroon have been provided in the references cited in the Study Site and Mosquito Sampling (Materials and Methods).

### Consent for publication

Not Applicable.

### Availability of data and materials

The dataset(s) supporting the conclusions of this article are available in the European Nucleotide Archive, with accession PRJEB51644, and secondary accession of ERP136291, for the RNA-seqe raw sequence reads. cDNA sequences of *GSTe2* and *CYP6Z2* were deposited in GenBank (accession numbers: ON169006 - ON169030 for *GSTe2* and ON169031 - ON169050 for *CYP6Z2*) and 5’-UTR DNA fragment sequences were deposited in GenBank (accession numbers: ON169051 - ON169140).

### Authors’ contributions

Conceived and designed by SSI and CSW. SSI carried out the molecular analyses, with support from AM, LMJM, EIP and HI. ANF and MMM, participated in field collection of mosquitoes in Nigeria, Chad and Cameroon. SSI carried out data analysis with the support of JH, GDW, SCN for the RNAseq component. SSI wrote the manuscript with inputs from CSW, JH and SCN. All authors contributed to corrections of the final draft and approved final version of the manuscript.

## Acknowledgements

Not Applicable

## Additional Files

**Additional File 1**: Supplementary text for methods and results.

**Additional File 2:** Common differentially expressed genes across the Sahel.

**Additional File 3:** Population genetics analyses.

**Additional File 4:** Genes within 2La, 2Rb and 2Rc inversion polymorphisms.

**Additional Tables:** Additional tables, S1-S6.

**Additional Figures:** Additional figures, S1-S11.

## Supplementary Figures

**Figure S1:**
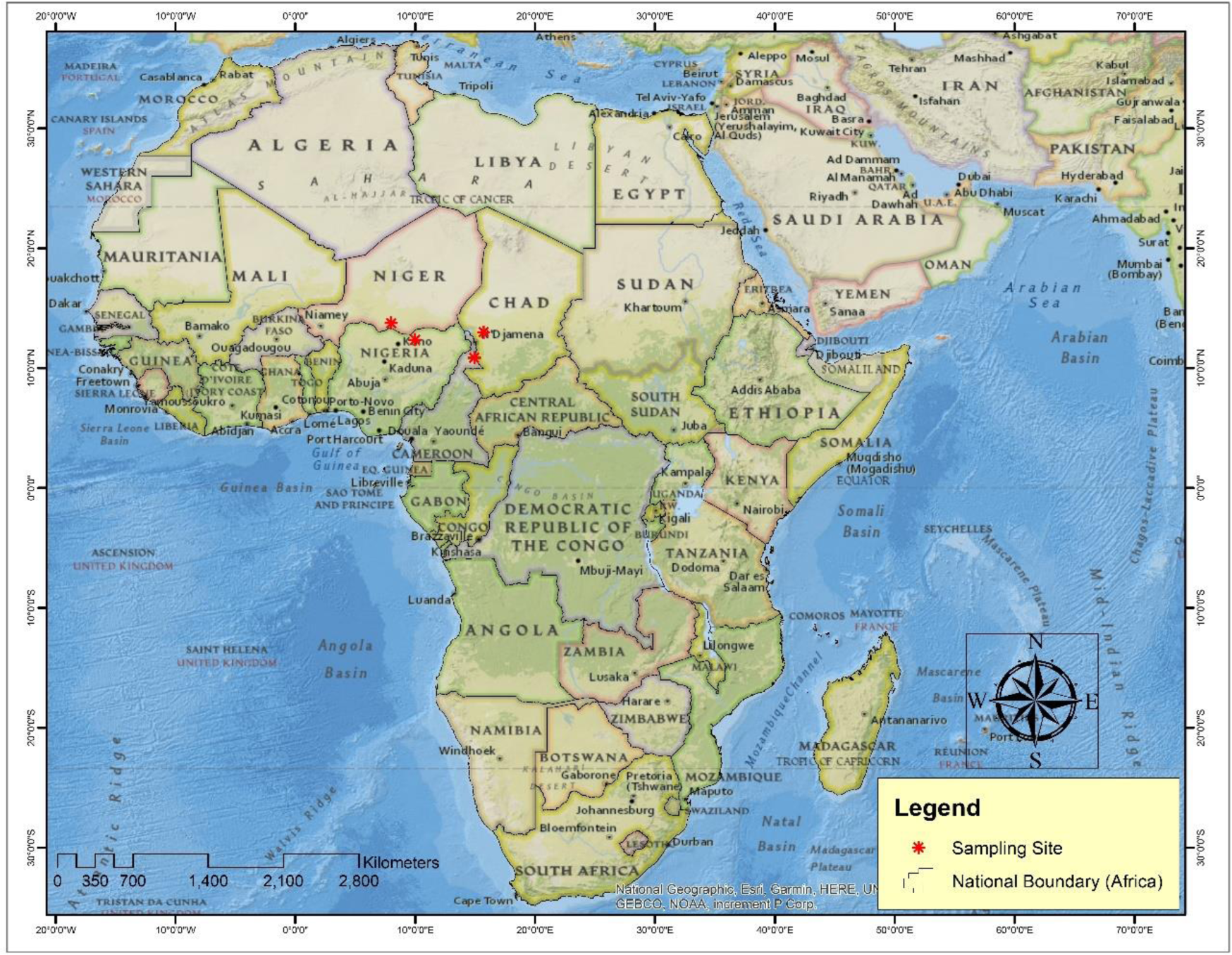
Sampling sites in the Sahel region of the four sampled countries.

**Figure S2:**
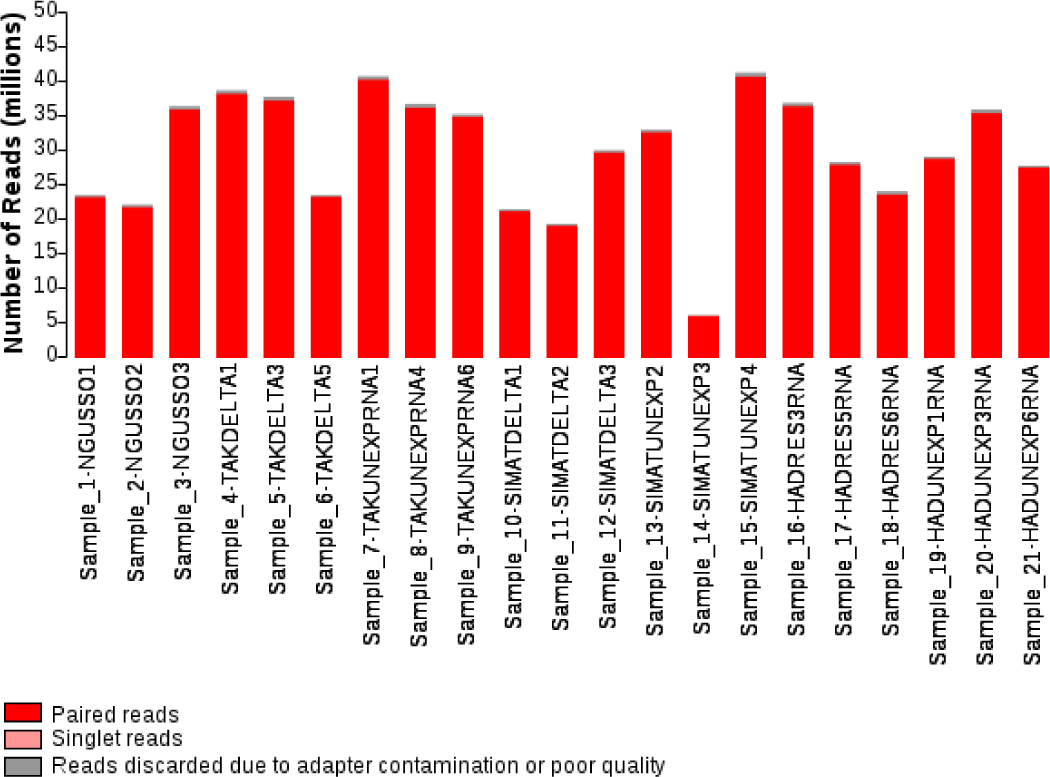
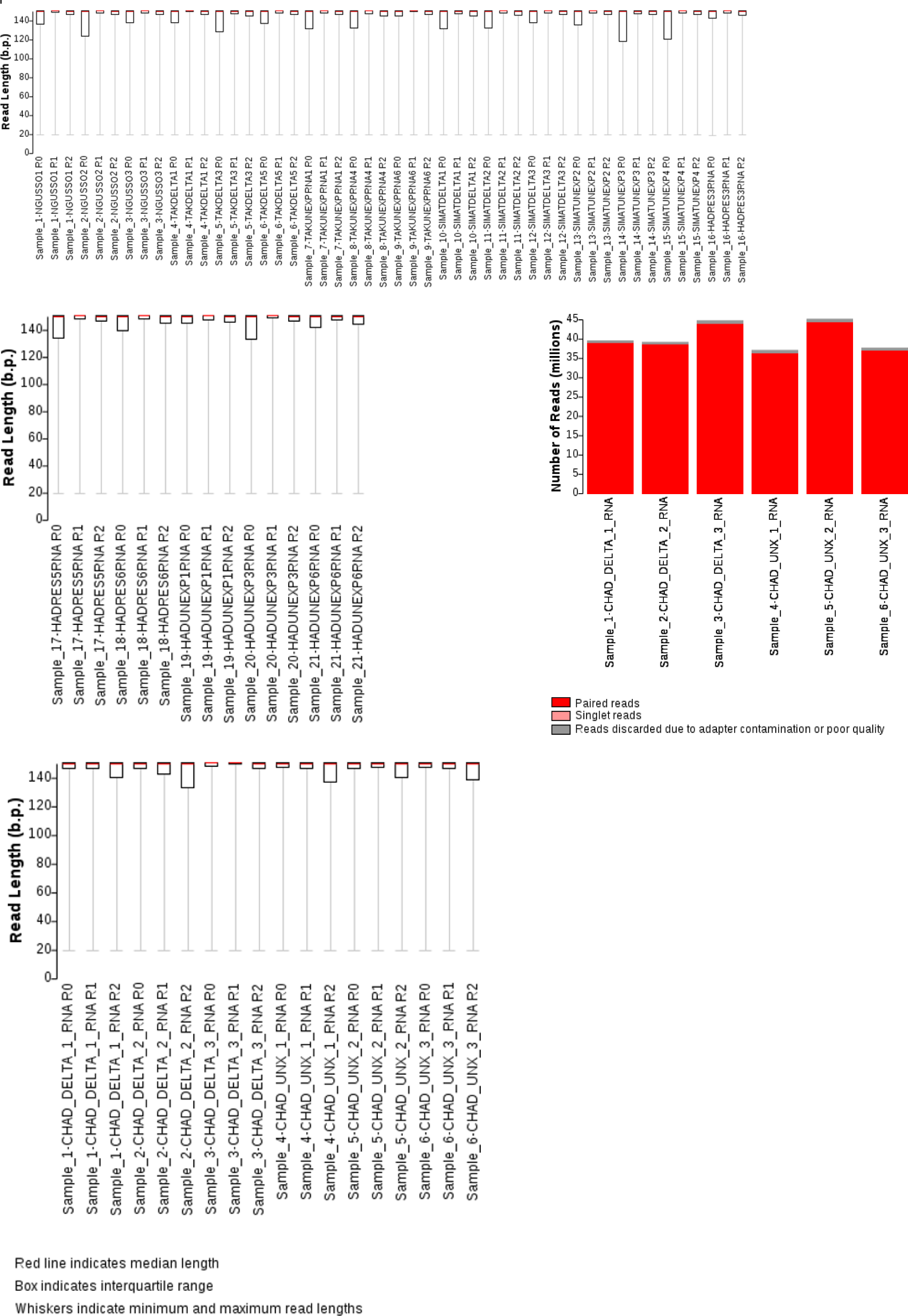
Summary of total number of reads for each sample and distribution of trimmed read length for forward (R1) and reverse (R2) reads and reads unpaired after trimming (R0).

**Figure S3:**
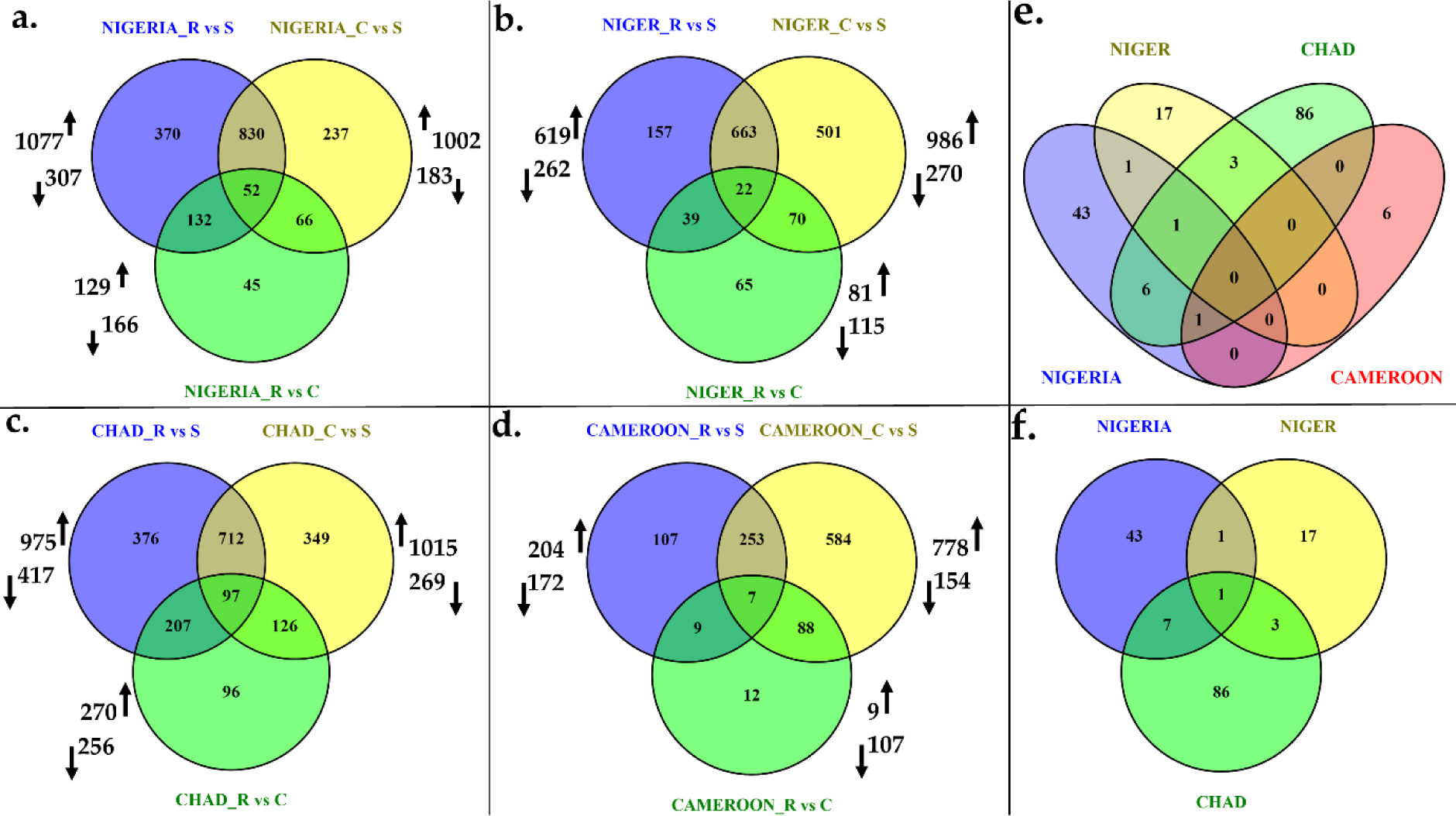
Venn diagrams summarizing the differentially expressed (DE) genes between resistant (R), unexposed (C) and susceptible (S) samples with a transcription ratio log_2_FC ≥ 1 in either direction, and a corrected p value *<* 0.05.

**Figure S4:**
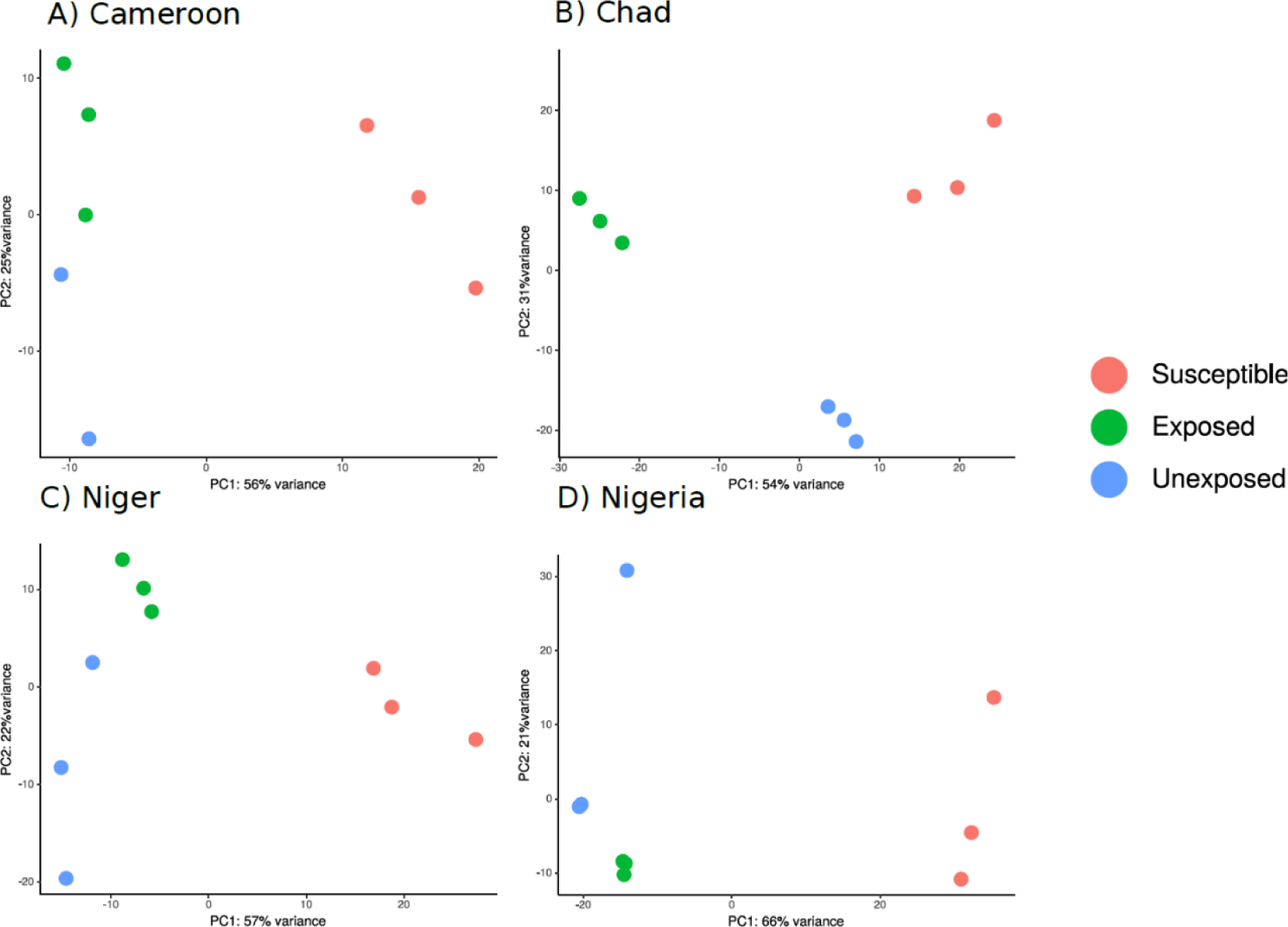
Principal component analysis of the 500 most variable genes from all experimental arms in the data from the four countries.

**Figure S5:**
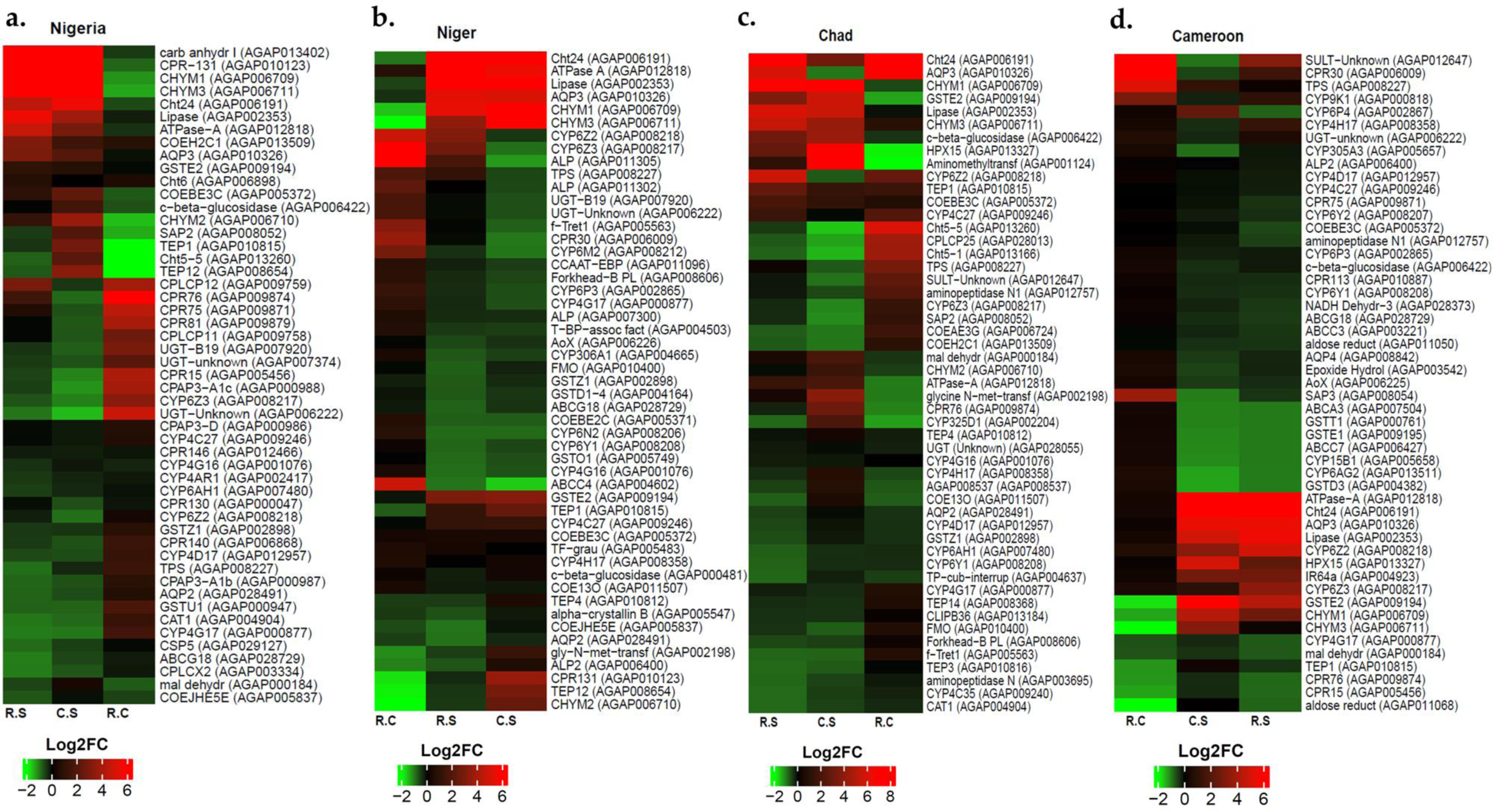
A heatmap showing the top 50 overexpressed genes in *An. coluzzii* populations from the Sahel region of each country. a. for Nigeria **b.**, **c.** and **d.** for Niger, Chad and Cameroon. Several genes including chymotrypsins, CYP450s, aquaporins, glutathione S-transferases and carboxylesterases are commonly upregulated across the Sahel of the four countries.

**Figure S6:**
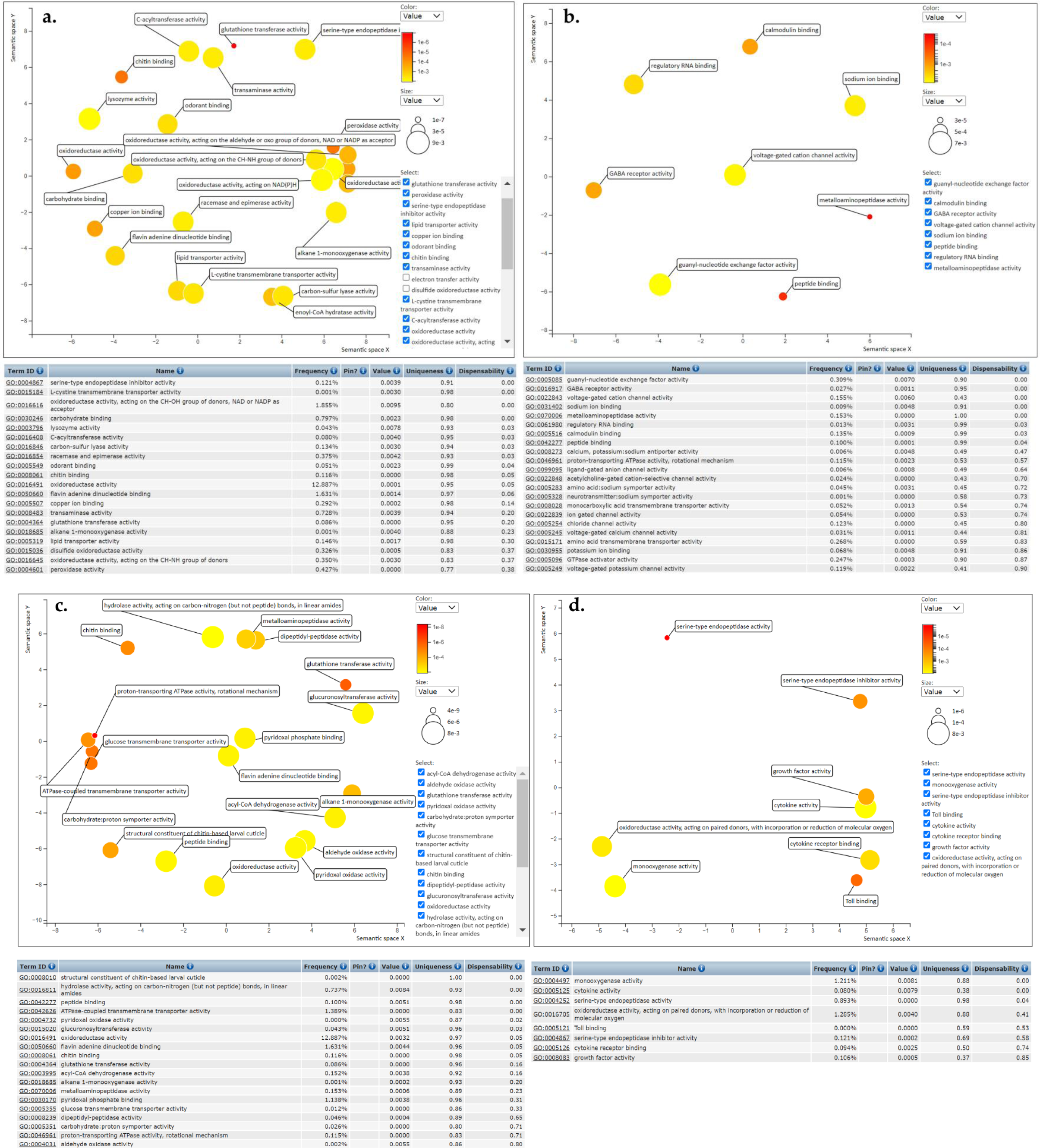
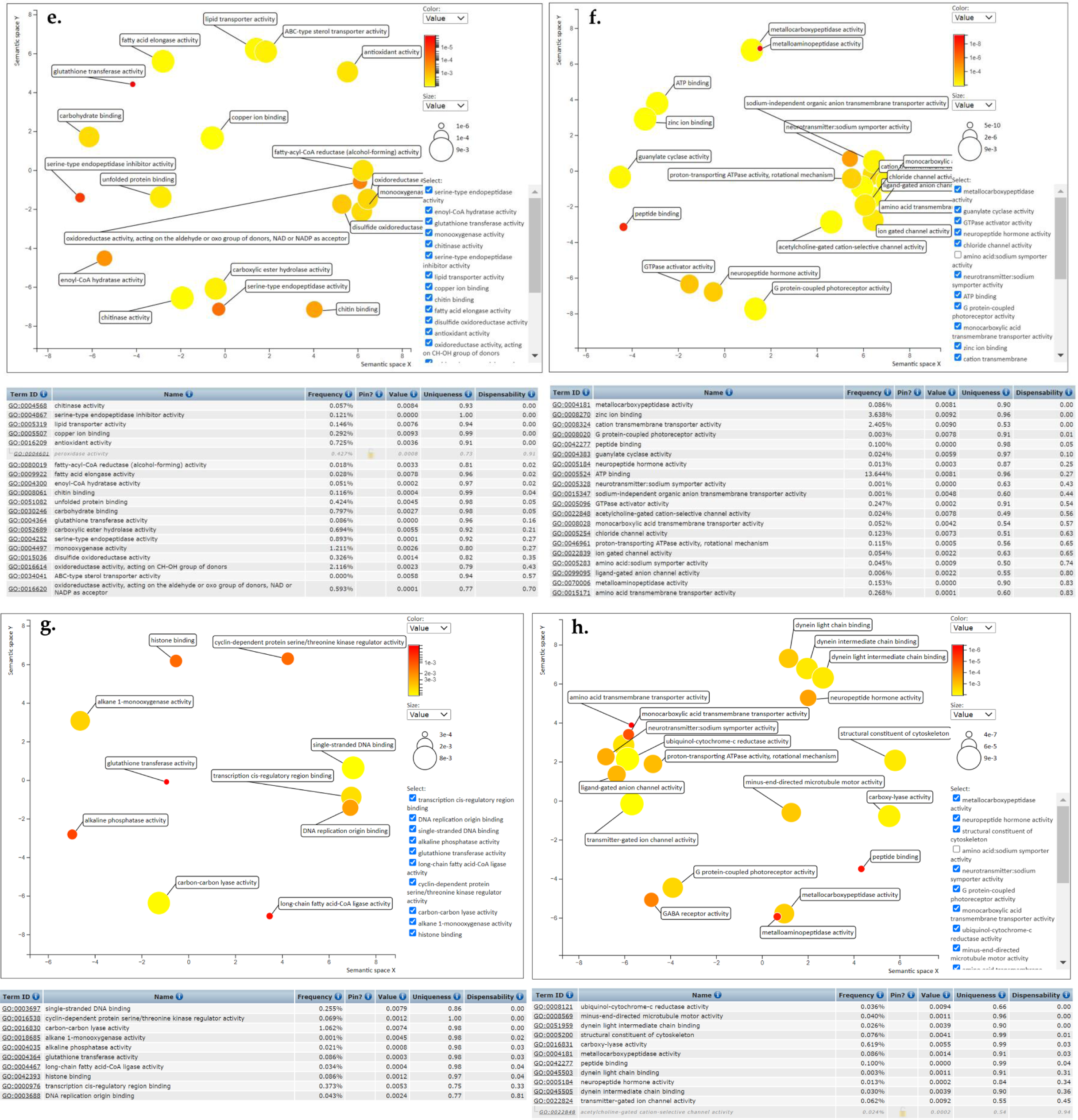
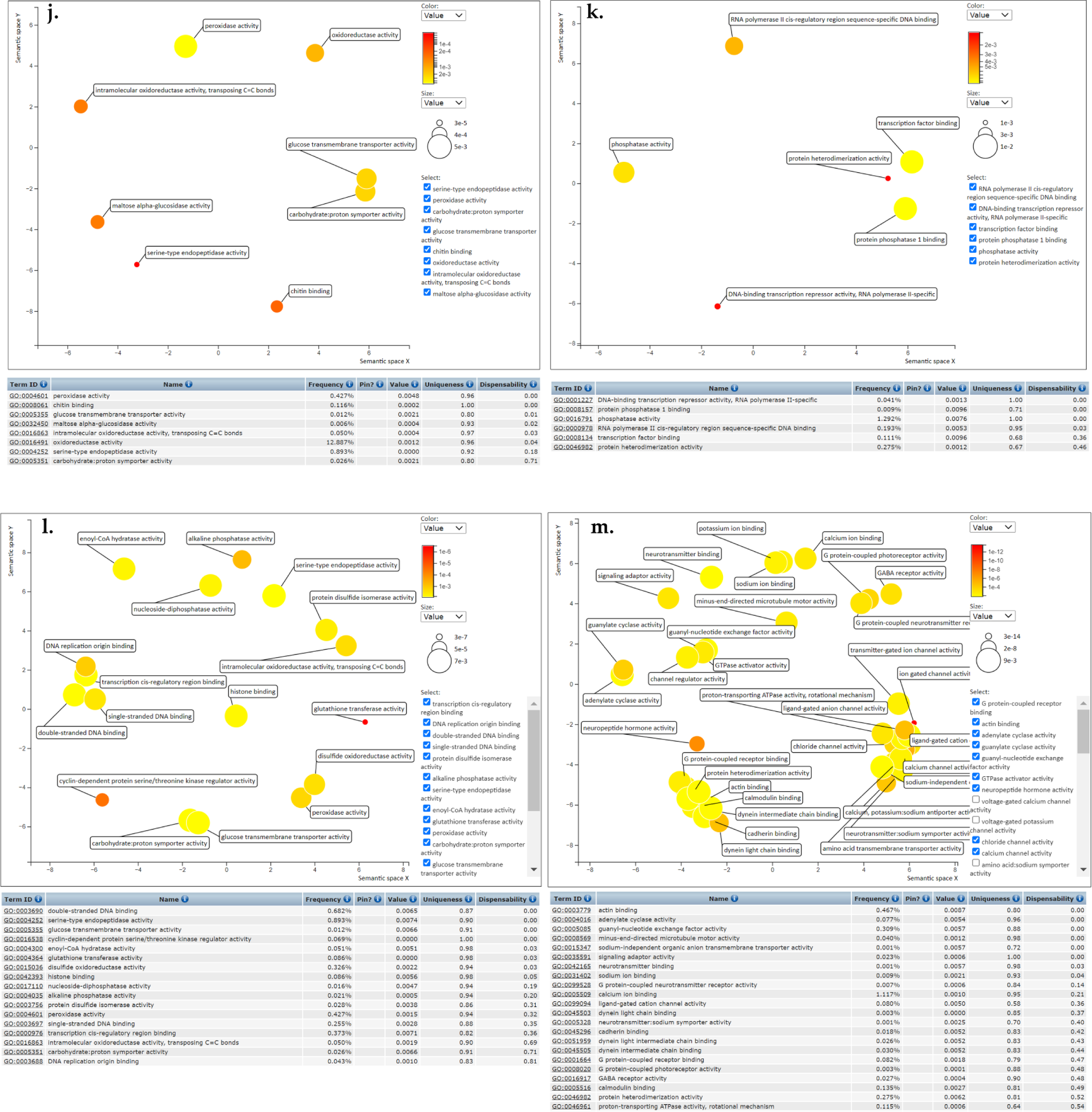
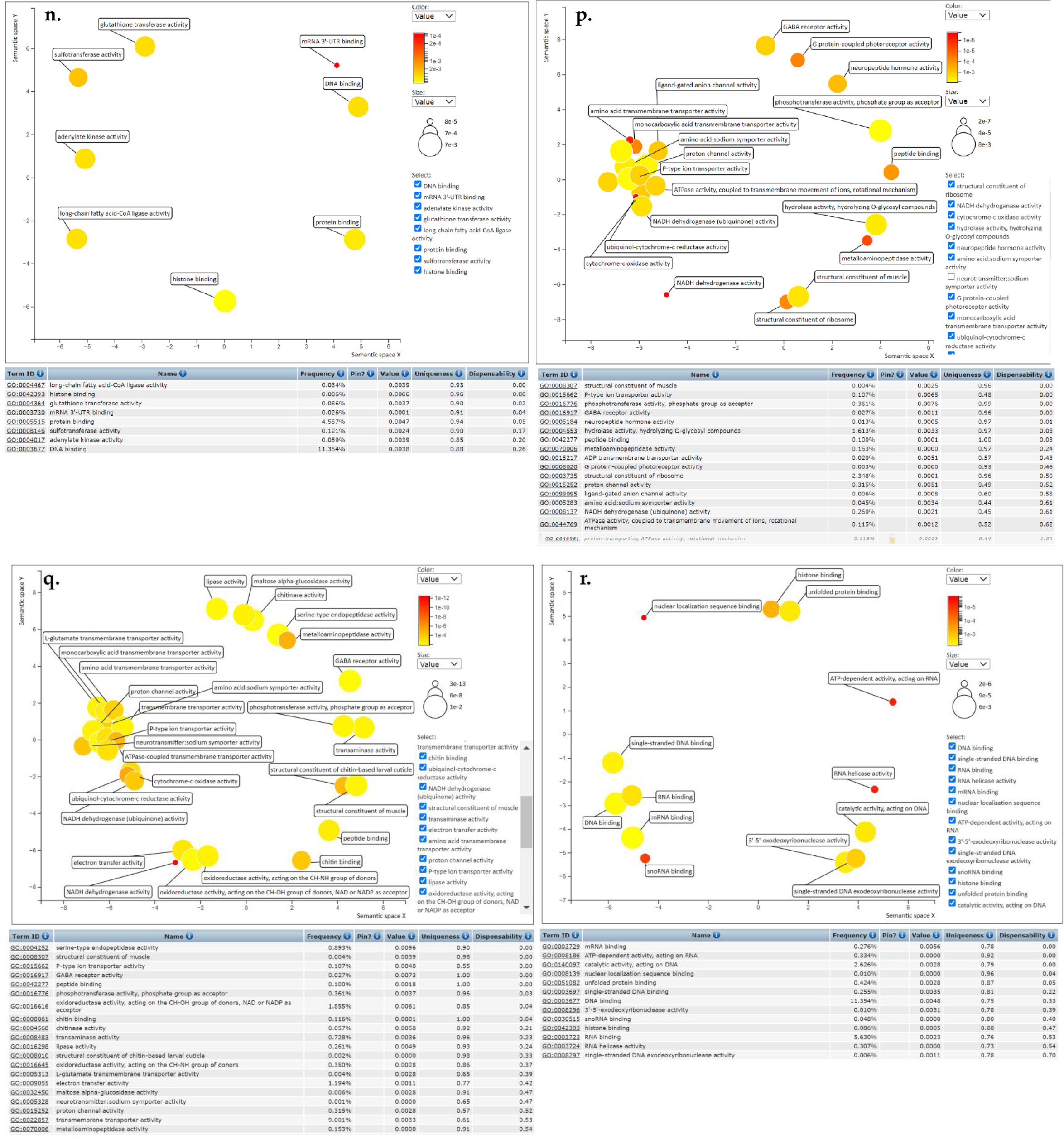
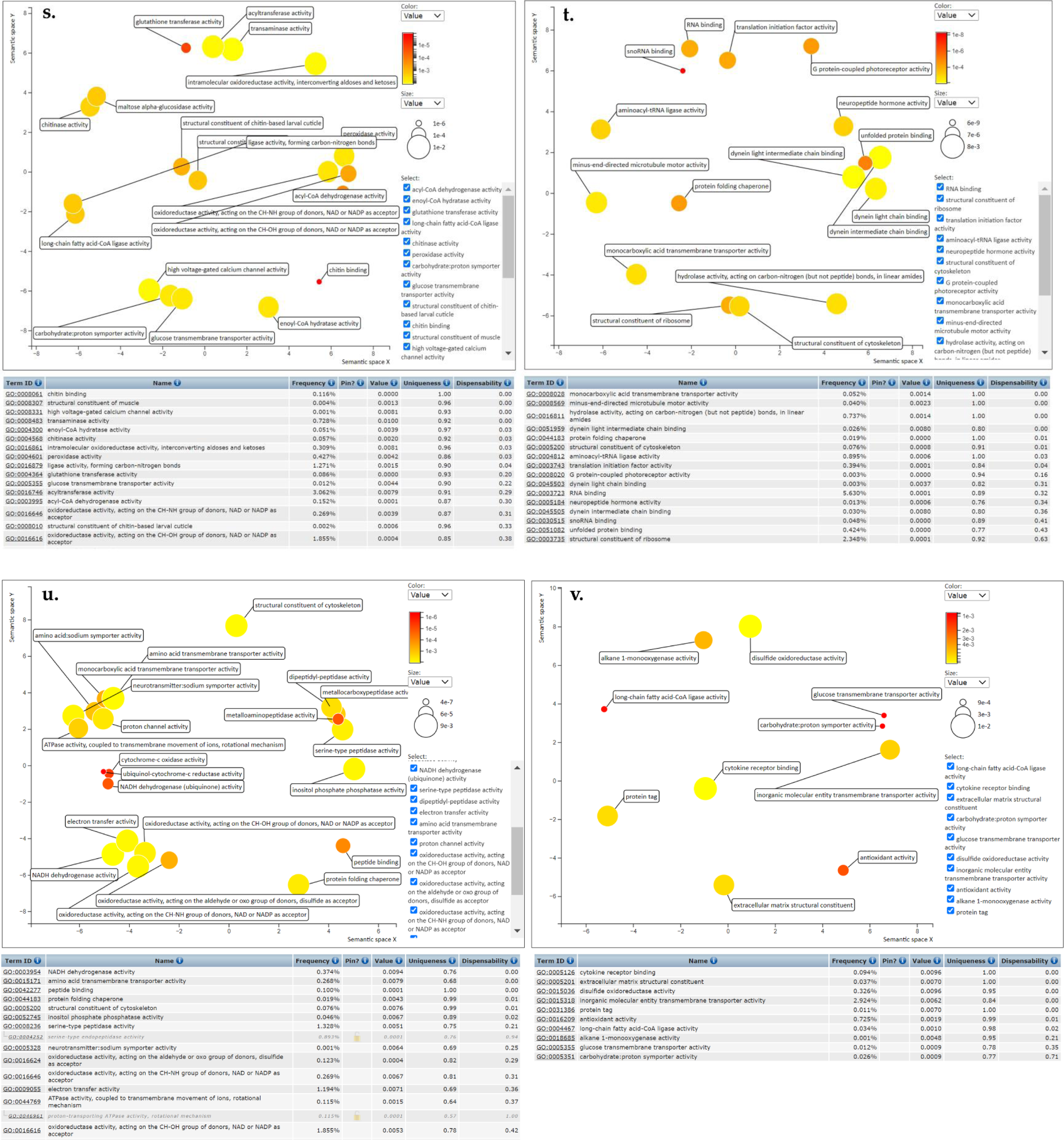
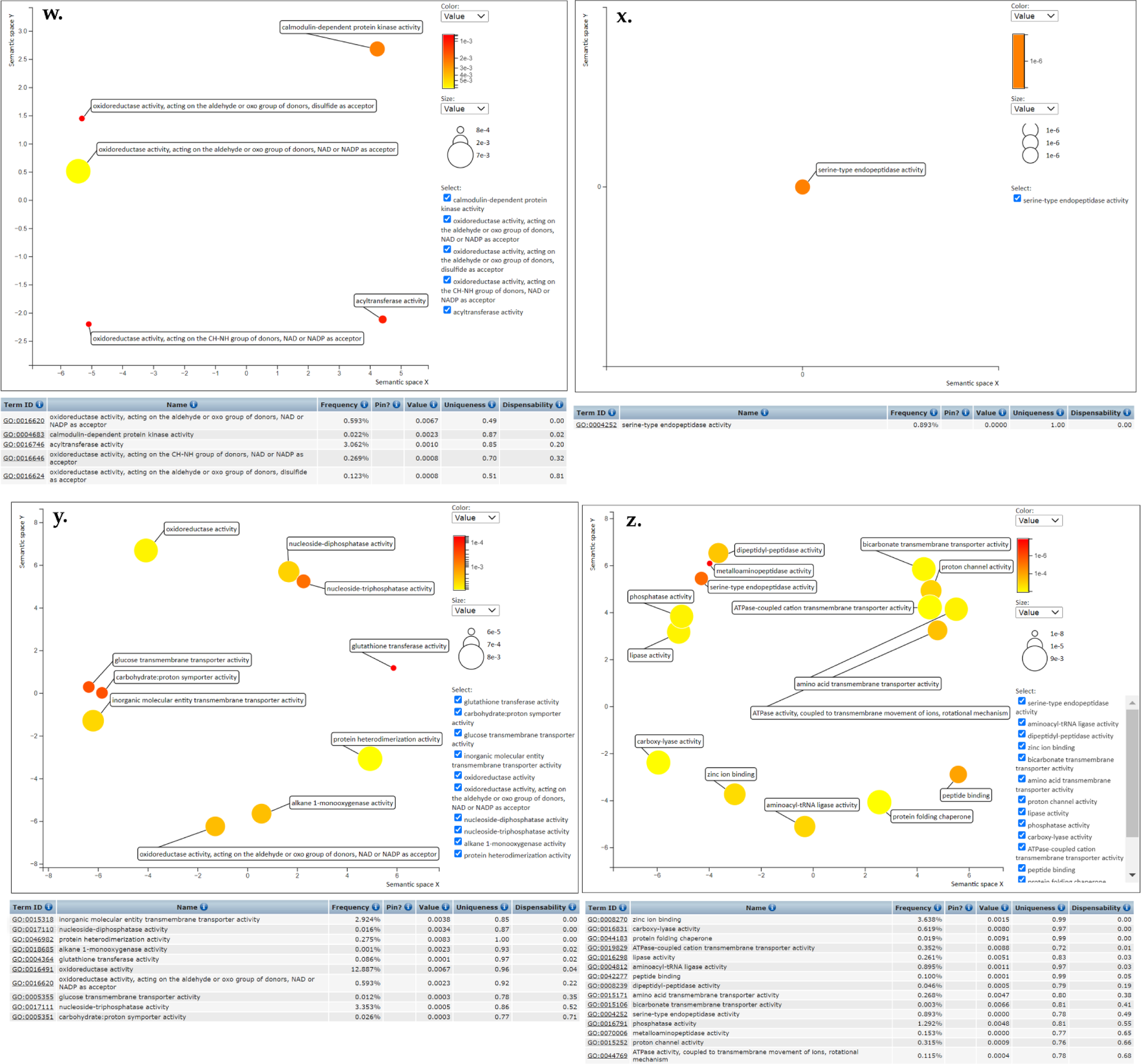
Revigo Scatter and Table Views of the GO terms over-represented in all comparisons (R vs S, R vs C and C vs S) for data from Nigeria, Niger, Chad and Cameroon.

**Figure S7:**
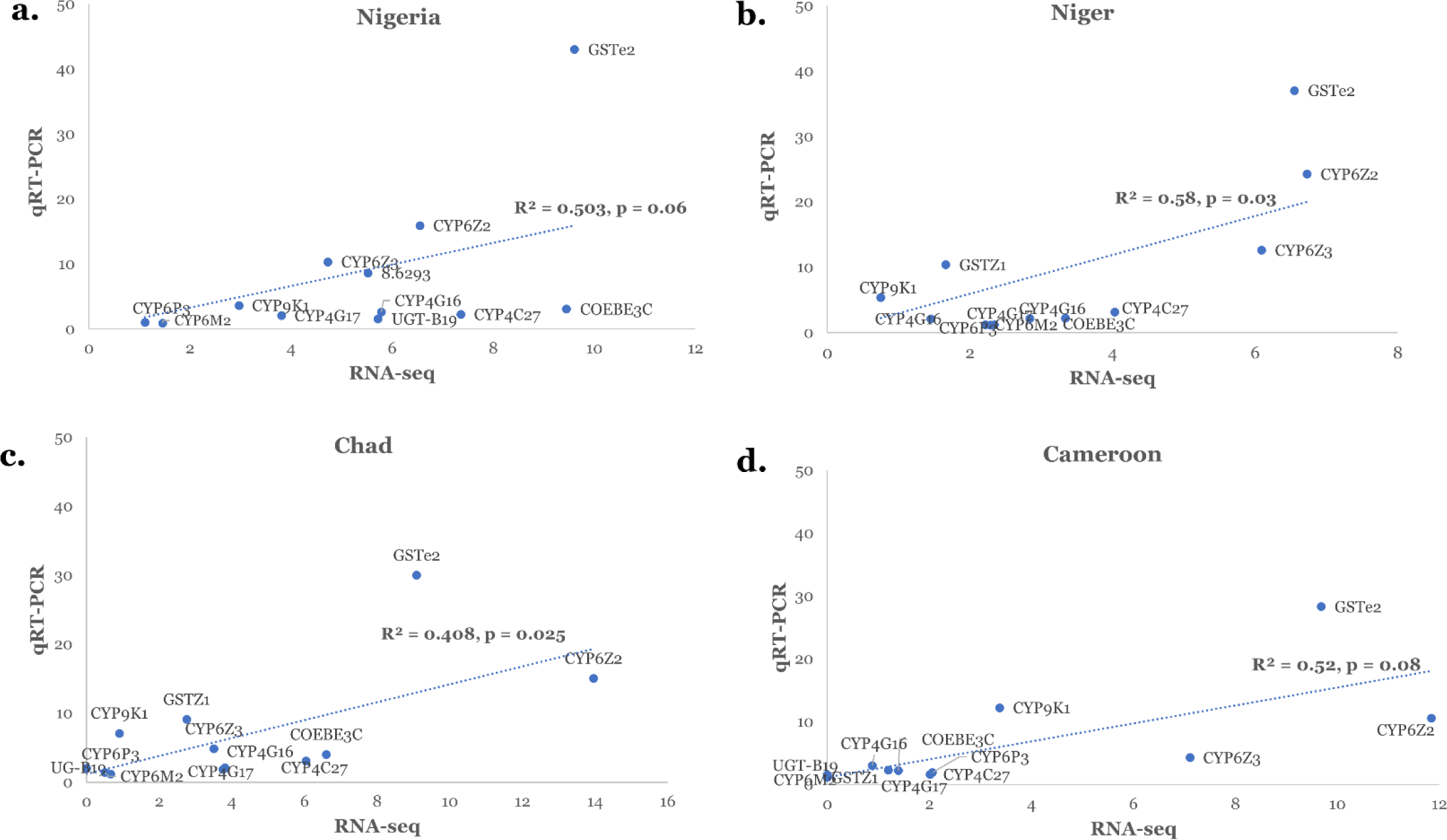
Validation of RNA-seq results using qRT-PCR analysis: correlation between RNA-seq fold changes and those obtained from qRT-PCR for 12 metabolic resistance genes.

**Figure S8:**
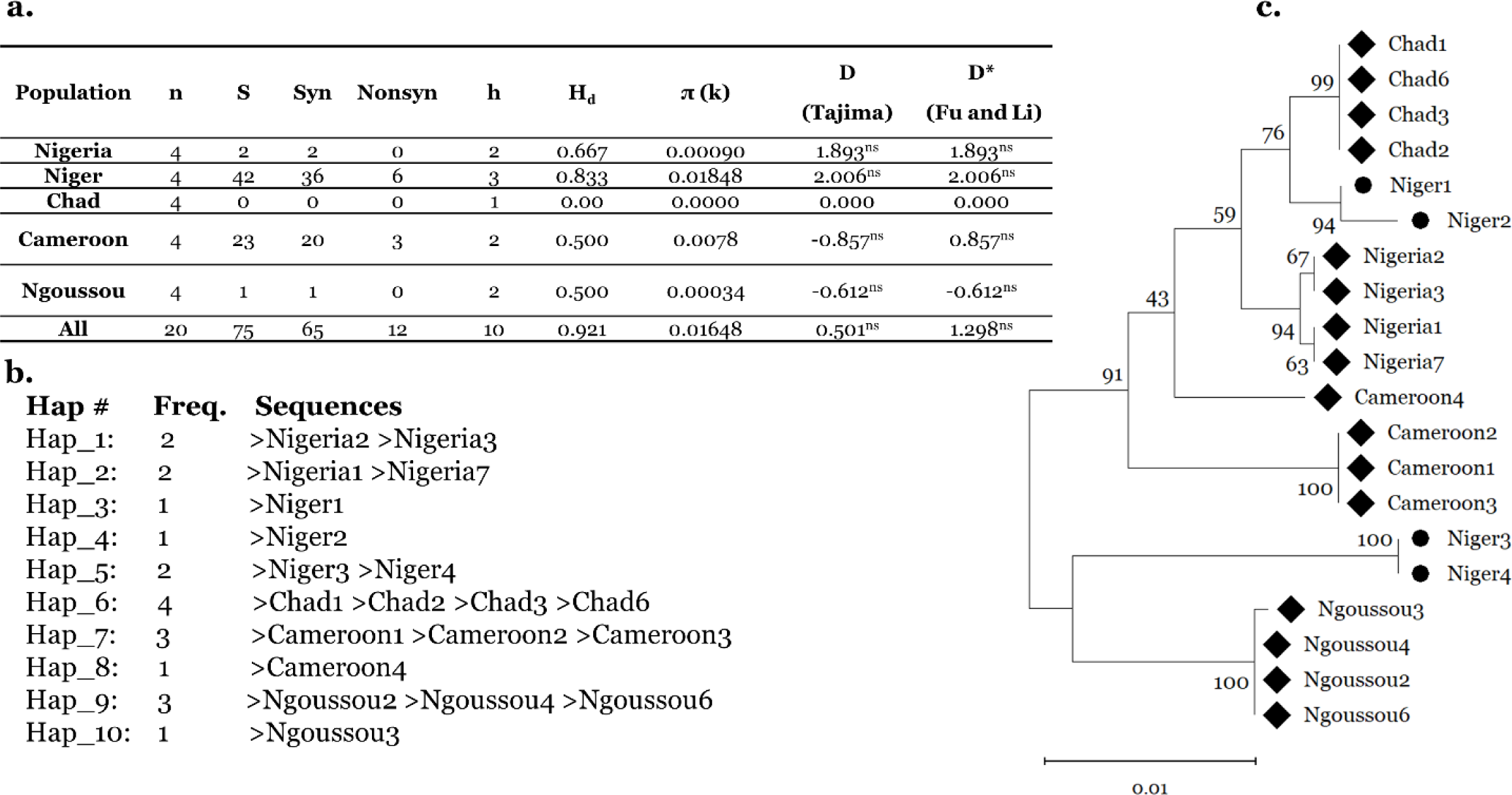
Polymorphism analysis of full length cDNA of *CYP6Z2*. **a.** Summary statistics for polymorphisms of *CYP6Z2* cDNA across the Sahel countries. n = number of sequences (n); S, number of polymorphic sites; h, haplotype; H_d_, haplotype diversity Syn, Synonymous mutations; Nonsyn, Non-synonymous mutations; π, nucleotide diversity (k= mean number of nucleotide differences); Tajima’s D and Fu and Li’s D statistics, ns, not significant. **b.** *CYP6Z2* cDNA haplotypes and its frequencies, **c.** a maximum likelihood phylogenetic tree of *CYP6Z2* cDNA sequences.

**Figure S9:**
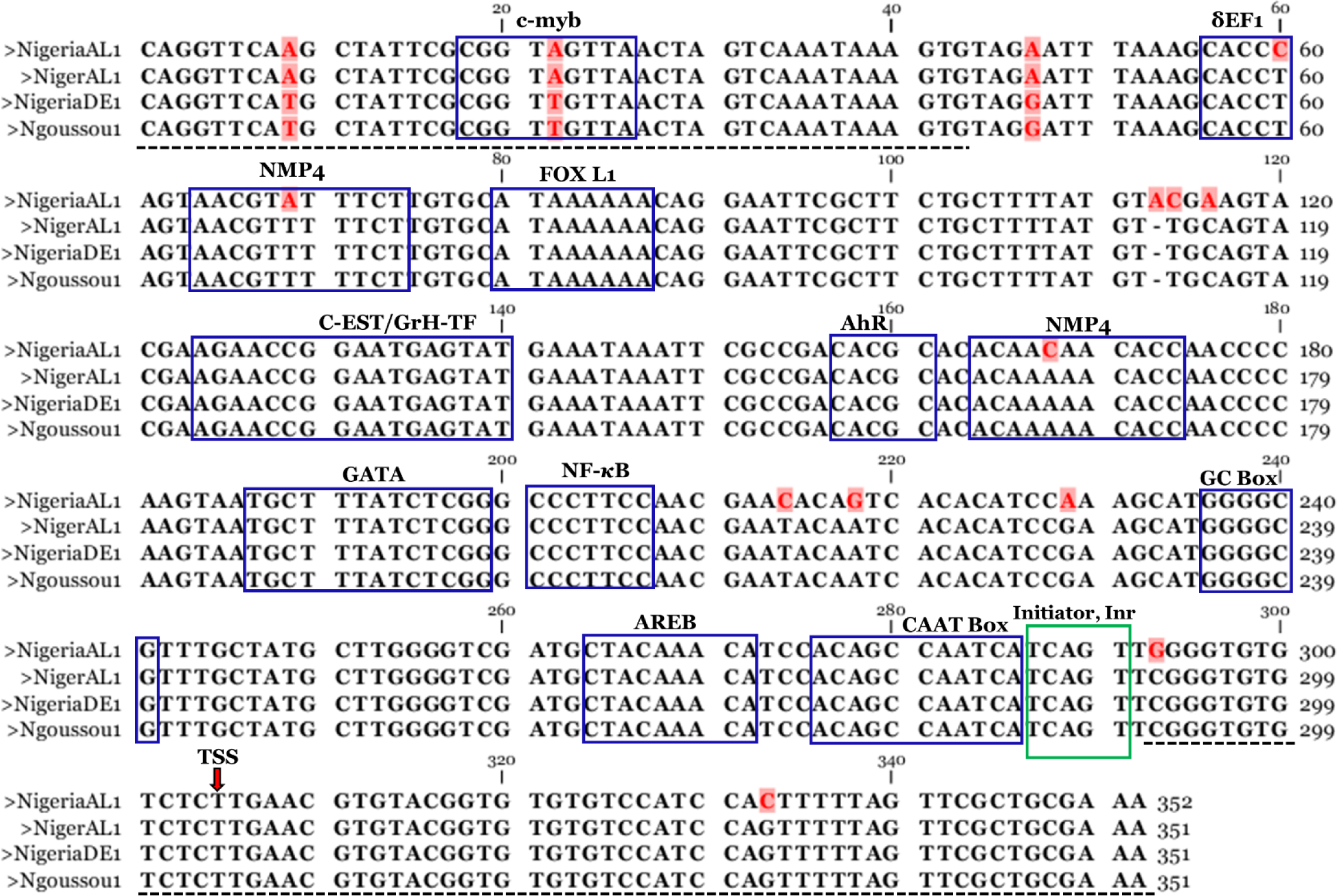
Comparative alignment of the *GSTe2* 5’-UTR fragments from the various haplotypes, showing putative transcription factors binding sites (blue boxes), and polymorphic positions (in red and highlighted in pink). The arthropod Inr consensus sequence is in green box, the transcription start site indicated with red arrow, and the 3’-UTR of *GSTe1* and 5’-UTR of *GSTe2* indicated with dashed lines.

**Figure S10:**
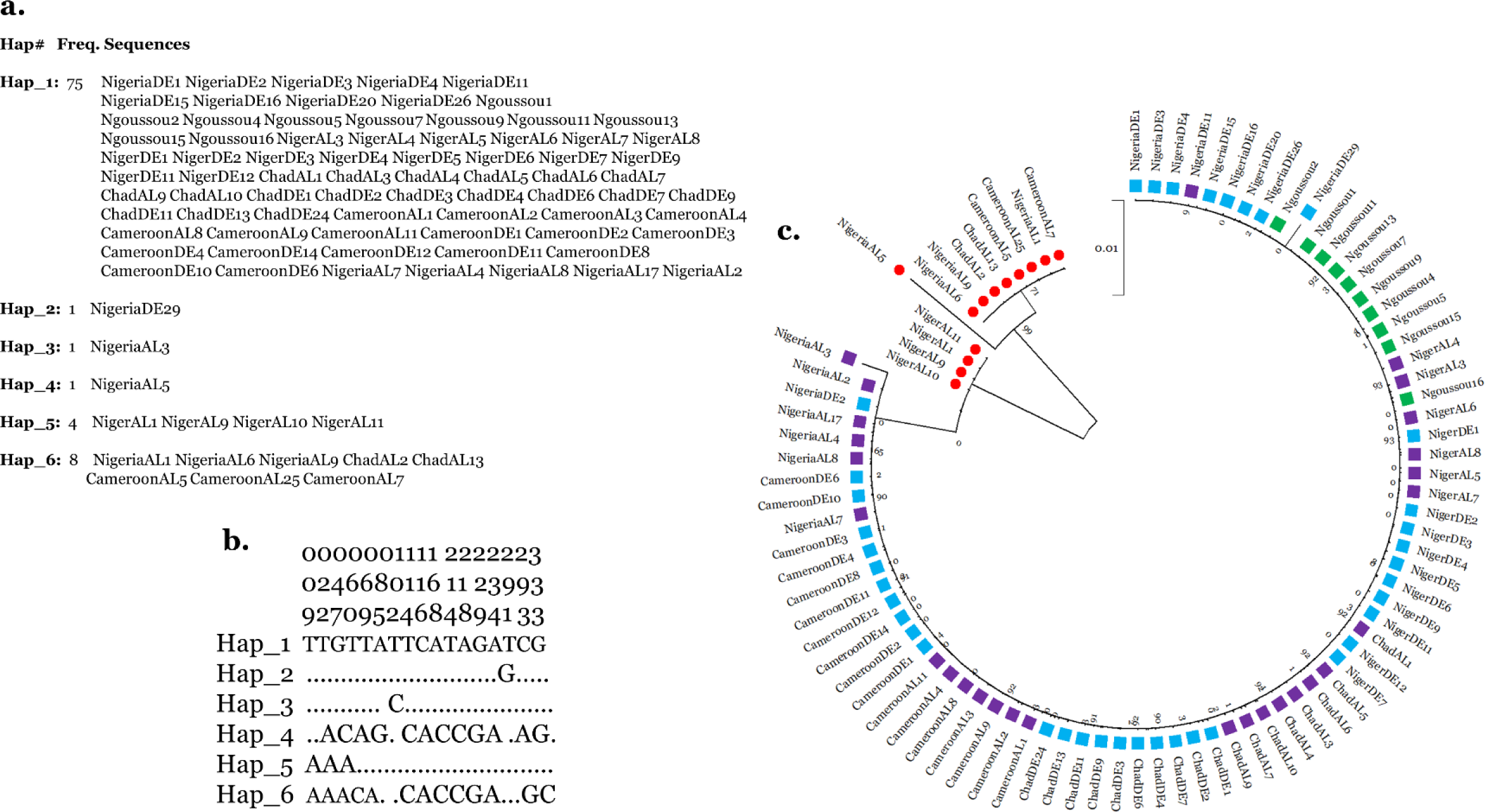
Genetic variability of the *GSTe2* 5’-UTR fragment from Nigeria, Niger, Chad, Cameroon and Ngoussou populations. **a.** the six haplotypes of the 5’-UTR with respective frequencies, **b.** polymorphic positions with numbers on top indicating the nucleotide position in the 351 bp fragment, **c.** maximum likelihood phylogenetic tree of the 90 sequences, showing the DDT-alive sequences with mutations forming a clade specific to its phenotype.

**Figure S11:**
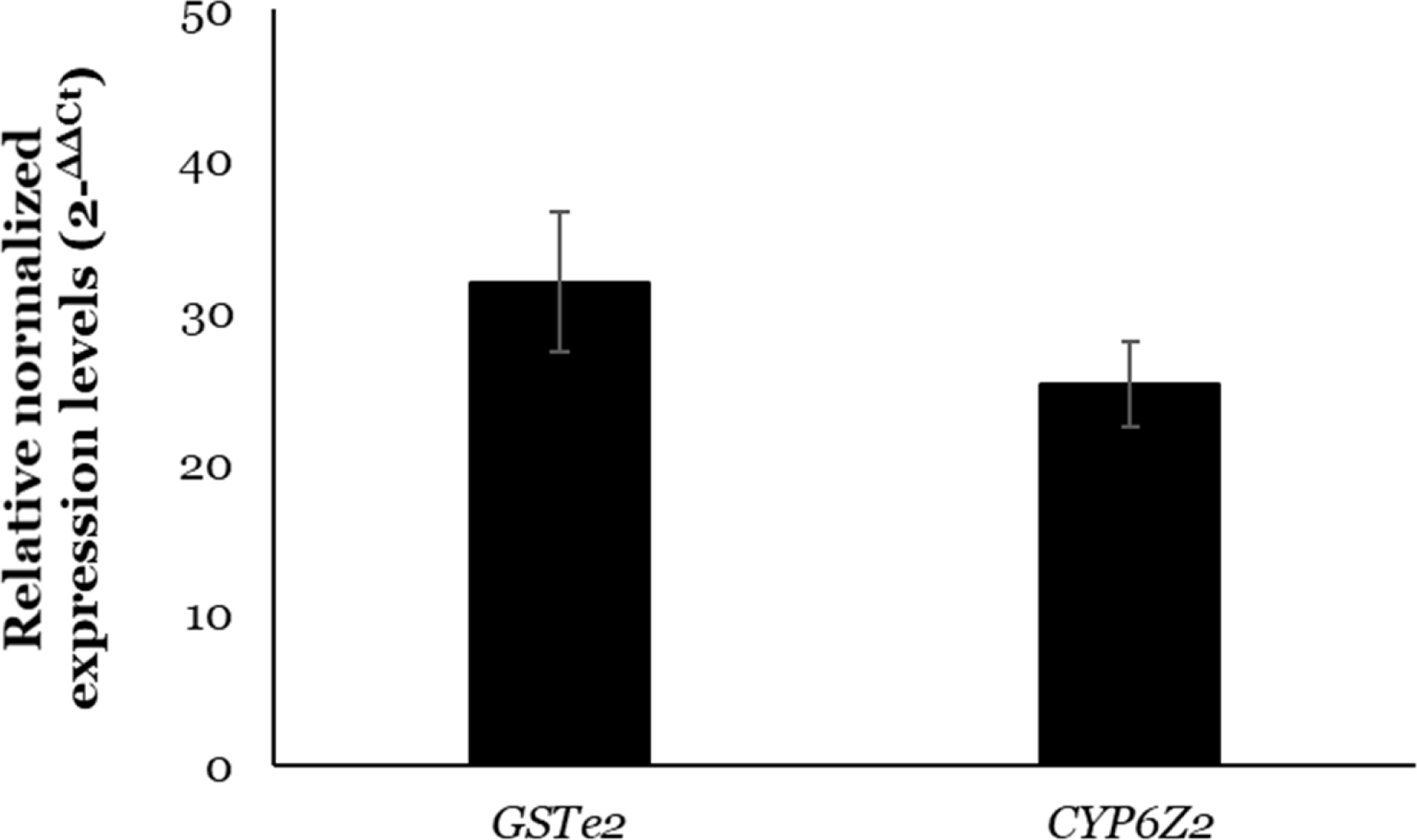
qRT-PCR validation of the overexpression of *GSTe2* and *CYP6Z2.* The relative expression of the *GSTe2* and *CYP6Z2* in the experimental transgenic *D. melanogaster* in relation to control flies. Data shown are the mean ± standard deviation (*n* = 6).

## Reference

1. Bhatt S, Weiss D, Cameron E, Bisanzio D, Mappin B, Dalrymple U, et al. The effect of malaria control on Plasmodium falciparum in Africa between 2000 and 2015. Nature. 2015;526(7572):207.

2. WHO. Global Technical Strategy for Malaria 2016-2030. World Health Organisation. . Geneva, Switzerland; 2015 May 2015. Report No.: ISBN 978 92 4 156499 1.

3. WHO. World Malaria Report. World Health Organization. . Geneva Switzerland; 2018. Report No.: Licence: CC BY-NC-SA 3.0 IGO Contract No.: ISBN 978-92-4-156565-3.

4. WHO. World Malaria Report. Geneva: Switzerland; 2021. Report No.: 978–92-4-004049-6 Contract No.: 978-92-4-004049-6

5. WHO. World Malaria Report 2019. Geneva, Switzerland; 2019. Report No.: 978-92-4-156572-1.

6. WHO. Update on the E-2020 initiative of 21 malaria-eliminating countries: report and country briefs. Geneva. Switzerland: World Health Organization; 2018. Report No.: WHO/CDS/GMP/2018.10.

7. Ibrahim SS, Mukhtar MM, Datti JA, Irving H, Kusimo MO, Tchapga W, et al. Temporal escalation of Pyrethroid Resistance in the major malaria vector Anopheles coluzzii from Sahelo-Sudanian Region of northern Nigeria. Scientific reports. 2019;9(1):7395.

8. Riveron JM, Huijben S, Tchapga W, Tchouakui M, Wondji MJ, Tchoupo M, et al. Escalation of Pyrethroid Resistance in the Malaria Vector Anopheles funestus Induces a Loss of Efficacy of Piperonyl Butoxide-Based Insecticide-Treated Nets in Mozambique. J Infect Dis. 2019;220(3):467–75.

9. Toe KH, N’Fale S, Dabire RK, Ranson H, Jones CM. The recent escalation in strength of pyrethroid resistance in Anopheles coluzzi in West Africa is linked to increased expression of multiple gene families. BMC Genomics. 2015;16:146.

10. Weedall GD, Mugenzi LMJ, Menze BD, Tchouakui M, Ibrahim SS, Amvongo-Adjia N, et al. A cytochrome P450 allele confers pyrethroid resistance on a major African malaria vector, reducing insecticide-treated bednet efficacy. Sci Transl Med. 2019;11(484).

11. Mugenzi LMJ, Menze BD, Tchouakui M, Wondji MJ, Irving H, Tchoupo M, et al. Cis-regulatory CYP6P9b P450 variants associated with loss of insecticide-treated bed net efficacy against Anopheles funestus. Nat Commun. 2019;10(1):4652.

12. Barnes KG, Weedall GD, Ndula M, Irving H, Mzihalowa T, Hemingway J, et al. Genomic Footprints of Selective Sweeps from Metabolic Resistance to Pyrethroids in African Malaria Vectors Are Driven by Scale up of Insecticide-Based Vector Control. PLoS Genet. 2017;13(2):e1006539.

13. Weedall GD, Riveron JM, Hearn J, Irving H, Kamdem C, Fouet C, et al. An Africa-wide genomic evolution of insecticide resistance in the malaria vector Anopheles funestus involves selective sweeps, copy number variations, gene conversion and transposons. PLoS Genet. 2020;16(6):e1008822.

14. Bruce-Chwatt LJ. Malaria in Nigeria. Bulletin of the World Health Organization. 1951;4(3):301–27.

15. Anopheles gambiae Genomes C, Data analysis g, Partner working g, Sample c-A, Burkina F, Cameroon, et al. Genetic diversity of the African malaria vector Anopheles gambiae. Nature. 2017;552(7683):96–100.

16. Muller P, Warr E, Stevenson BJ, Pignatelli PM, Morgan JC, Steven A, et al. Field-caught permethrin-resistant Anopheles gambiae overexpress CYP6P3, a P450 that metabolises pyrethroids. PLoS Genet. 2008;4(11):e1000286.

17. Yunta C, Hemmings K, Stevenson B, Koekemoer LL, Matambo T, Pignatelli P, et al. Cross-resistance profiles of malaria mosquito P450s associated with pyrethroid resistance against WHO insecticides. Pestic Biochem Physiol. 2019;161:61–7.

18. Stevenson BJ, Bibby J, Pignatelli P, Muangnoicharoen S, O’Neill PM, Lian LY, et al. Cytochrome P450 6M2 from the malaria vector Anopheles gambiae metabolizes pyrethroids: Sequential metabolism of deltamethrin revealed. Insect biochemistry and molecular biology. 2011;41(7):492–502.

19. Mitchell SN, Stevenson BJ, Müller P, Wilding CS, Egyir-Yawson A, Field SG, et al. Identification and validation of a gene causing cross-resistance between insecticide classes in Anopheles gambiae from Ghana. Proceedings of the National Academy of Sciences. 2012;109(16):6147–52.

20. Mitchell SN, Rigden DJ, Dowd AJ, Lu F, Wilding CS, Weetman D, et al. Metabolic and target-site mechanisms combine to confer strong DDT resistance in Anopheles gambiae. PLoS One. 2014;9(3):e92662.

21. Lucas ER, Rockett KA, Lynd A, Essandoh J, Grisales N, Kemei B, et al. A high throughput multi-locus insecticide resistance marker panel for tracking resistance emergence and spread in Anopheles gambiae. Sci Rep. 2019;9(1):13335.

22. Riveron JM, Yunta C, Ibrahim SS, Djouaka R, Irving H, Menze BD, et al. A single mutation in the GSTe2 gene allows tracking of metabolically based insecticide resistance in a major malaria vector. Genome Biol. 2014;15(2):R27.

23. Menze BD, Kouamo MF, Wondji MJ, Tchapga W, Tchoupo M, Kusimo MO, et al. An Experimental Hut Evaluation of PBO-Based and Pyrethroid-Only Nets against the Malaria Vector Anopheles funestus Reveals a Loss of Bed Nets Efficacy Associated with GSTe2 Metabolic Resistance. Genes (Basel). 2020;11(2).

24. Coluzzi M, Sabatini A, Petrarca V, Di Deco MA. Chromosomal differentiation and adaptation to human environments in the Anopheles gambiae complex. Trans R Soc Trop Med Hyg. 1979;73(5):483–97.

25. Cairns M, Roca-Feltrer A, Garske T, Wilson AL, Diallo D, Milligan PJ, et al. Estimating the potential public health impact of seasonal malaria chemoprevention in African children. Nat Commun. 2012;3:881.

26. Ibrahim SS, Mukhtar MM, Irving H, Labbo R, Kusimo MO, Mahamadou I, et al. High Plasmodium infection and multiple insecticide resistance in a major malaria vector Anopheles coluzzii from Sahel of Niger Republic. Malaria journal. 2019;18(1):181.

27. Fadel AN, Ibrahim SS, Tchouakui M, Terence E, Wondji MJ, Tchoupo M, et al. A combination of metabolic resistance and high frequency of the 1014F kdr mutation is driving pyrethroid resistance in Anopheles coluzzii population from Guinea savanna of Cameroon. Parasit Vectors. 2019;12(1):263.

28. Ibrahim SS, Fadel AN, Tchouakui M, Terence E, Wondji MJ, Tchoupo M, et al. High insecticide resistance in the major malaria vector Anopheles coluzzii in Chad Republic. Infect Dis Poverty. 2019;8(1):100.

29. Mitchell SN, Stevenson BJ, Muller P, Wilding CS, Egyir-Yawson A, Field SG, et al. Identification and validation of a gene causing cross-resistance between insecticide classes in Anopheles gambiae from Ghana. Proceedings of the National Academy of Sciences of the United States of America. 2012;109(16):6147–52.

30. Martin M. Cutadapt removes adapter sequences from high-throughput sequencing reads. EMBnet journal. 2011;17(1):10–2.

31. Joshi N, Fass J. Sickle: A sliding-window, adaptive, quality-based trimming tool for FastQ files (Version 1.33)[Software]. 2011.

32. Aronesty E. ea-utils: Command-line tools for processing biological sequencing data. Durham, NC; 2011.

33. Love MI, Huber, W., & Anders, S. Moderated estimation of fold change and dispersion for RNA-seq data with DESeq2. Genome Biology. 2014;15(550).

34. Blighe K, S. Rana, and M. Lewis. . “EnhancedVolcano: Publication-ready volcano plots with enhanced colouring and labeling.”. 2018.

35. Gu Z, Eils R, Schlesner M. Complex heatmaps reveal patterns and correlations in multidimensional genomic data. Bioinformatics. 2016;32(18):2847–9.

36. Alexa A. RJ. topGO: Enrichment Analysis for Gene Ontology. R package version 2.46.0. 2021 [

37. Supek F, Bosnjak M, Skunca N, Smuc T. REVIGO summarizes and visualizes long lists of gene ontology terms. PLoS One. 2011;6(7):e21800.

38. Riveron JM, Ibrahim SS, Chanda E, Mzilahowa T, Cuamba N, Irving H, et al. The highly polymorphic CYP6M7 cytochrome P450 gene partners with the directionally selected CYP6P9a and CYP6P9b genes to expand the pyrethroid resistance front in the malaria vector Anopheles funestus in Africa. BMC Genomics. 2014;15(1):817.

39. Schmittgen TD, Livak KJ. Analyzing real-time PCR data by the comparative C(T) method. Nature protocols. 2008;3(6):1101–8.

40. Nagi CS, Oruni A, Weetman D, Donnelly J. M. RNA-Seq-Pop: Exploiting the sequence in RNA-Seq - a Snakemake workflow reveals patterns of insecticide resistance in the malaria vector Anopheles gambiae. 2022;2022.06.17.493894.

41. Hudson RR, Slatkin M, Maddison WP. Estimation of levels of gene flow from DNA sequence data. Genetics. 1992;132(2):583–9.

42. Yi X, Liang Y, Huerta-Sanchez E, Jin X, Cuo ZX, Pool JE, et al. Sequencing of 50 human exomes reveals adaptation to high altitude. Science. 2010;329(5987):75–8.

43. Alistair Miles MFR, Peter Ralph, Nick Harding, Rahul Pisupati, and Summer Rae. cggh/scikit-allel: A Python package for exploring and analysing genetic variation data 2020 [cggh/scikit-allel: v1.3.1 (v.3.1). Zenodo.].

44. Li H, Handsaker B, Wysoker A, Fennell T, Ruan J, Homer N, et al. The Sequence Alignment/Map format and SAMtools. Bioinformatics. 2009;25(16):2078–9.

45. Love RR, Redmond SN, Pombi M, Caputo B, Petrarca V, Della Torre A, et al. In Silico Karyotyping of Chromosomally Polymorphic Malaria Mosquitoes in the Anopheles gambiae Complex. G3 (Bethesda). 2019;9(10):3249–62.

46. Hall TA, editor BioEdit: a user-friendly biological sequence alignment editor and analysis program for Windows 95/98/NT. Nucleic acids symposium series; 1999.

47. Kumar S, Stecher G, Li M, Knyaz C, Tamura K. MEGA X: Molecular Evolutionary Genetics Analysis across Computing Platforms. Molecular biology and evolution. 2018;35(6):1547–9.

48. Rozas J, Ferrer-Mata A, Sanchez-DelBarrio JC, Guirao-Rico S, Librado P, Ramos-Onsins SE, et al. DnaSP 6: DNA Sequence Polymorphism Analysis of Large Data Sets. Molecular biology and evolution. 2017;34(12):3299–302.

49. Cartharius K, Frech K, Grote K, Klocke B, Haltmeier M, Klingenhoff A, et al. MatInspector and beyond: promoter analysis based on transcription factor binding sites. Bioinformatics. 2005;21(13):2933–42.

50. Lynd A, Lycett GJ. Optimization of the Gal4-UAS system in an Anopheles gambiae cell line. Insect Mol Biol. 2011;20(5):599–608.

51. Ibrahim SS, Riveron JM, Bibby J, Irving H, Yunta C, Paine MJI, et al. Allelic Variation of Cytochrome P450s Drives Resistance to Bednet Insecticides in a Major Malaria Vector. PLoS Genet. 2015;11(10):e1005618.

52. Riveron JM, Tchouakui M., Mugenzi L., Menze B. D., Mu-Chun C. and Wondji C. S. Insecticide Resistance in Malaria Vectors: An Update at a Global Scale. In: S. Manguin VD, editor. Towards Malaria Elimination - A Leap Forward. London: IntechOpen; 2018. p. 452.

53. Shiff C. Integrated approach to malaria control. Clinical microbiology reviews. 2002;15(2):278–93.

54. Lindsay S, Parson, L. and Thomas, C. . Mapping the range and relative abundance of the two principal African malaria vectors, Anopheles gambiae sensu stricto and An. arabiensis, using climate data. Proceedings of the Royal Society of London B: Biological Science. 1998; 265((1399)): 847–54.

55. Coetzee M, Craig, M. and Le Sueur, D. . Distribution of African malaria mosquitoes belonging to the Anopheles gambiae complex. Parasitology today. 2000;16(2):74–7.

56. Ranson H, Abdallah H, Badolo A, Guelbeogo WM, Kerah-Hinzoumbe C, Yangalbe-Kalnone E, et al. Insecticide resistance in Anopheles gambiae: data from the first year of a multi-country study highlight the extent of the problem. Malar J. 2009;8:299.

57. Ibrahim SS, Mukhtar MM, Muhammad A, Wondji CS. 2La Paracentric Chromosomal Inversion and Overexpressed Metabolic Genes Enhance Thermotolerance and Pyrethroid Resistance in the Major Malaria Vector Anopheles gambiae. Biology (Basel). 2021;10(6).

58. Gimonneau G, Bouyer J, Morand S, Besansky NJ, Diabate A, Simard F. A behavioral mechanism underlying ecological divergence in the malaria mosquito Anopheles gambiae. Behav Ecol. 2010;21(5):1087–92.

59. Dao A, Yaro AS, Diallo M, Timbine S, Huestis DL, Kassogue Y, et al. Signatures of aestivation and migration in Sahelian malaria mosquito populations. Nature. 2014;516(7531):387–90.

60. Lehmann T, Weetman D, Huestis DL, Yaro AS, Kassogue Y, Diallo M, et al. Tracing the origin of the early wet-season Anopheles coluzzii in the Sahel. Evol Appl. 2017;10(7):704–17.

61. Adolfi A, Poulton B, Anthousi A, Macilwee S, Ranson H, Lycett GJ. Functional genetic validation of key genes conferring insecticide resistance in the major African malaria vector, Anopheles gambiae. Proceedings of the National Academy of Sciences of the United States of America. 2019;116(51):25764–72.

62. Tchouakui M, Riveron JM, Djonabaye D, Tchapga W, Irving H, Soh Takam P, et al. Fitness Costs of the Glutathione S-Transferase Epsilon 2 (L119F-GSTe2) Mediated Metabolic Resistance to Insecticides in the Major African Malaria Vector Anopheles Funestus. Genes (Basel). 2018;9(12).

63. Chiu TL, Wen Z, Rupasinghe SG, Schuler MA. Comparative molecular modeling of Anopheles gambiae CYP6Z1, a mosquito P450 capable of metabolizing DDT. Proceedings of the National Academy of Sciences of the United States of America. 2008;105(26):8855–60.

64. Yunta C, Grisales N, Nasz S, Hemmings K, Pignatelli P, Voice M, et al. Pyriproxyfen is metabolized by P450s associated with pyrethroid resistance in An. gambiae. Insect biochemistry and molecular biology. 2016;78:50–7.

65. Lees RS, Ismail HM, Logan RAE, Malone D, Davies R, Anthousi A, et al. New insecticide screening platforms indicate that Mitochondrial Complex I inhibitors are susceptible to cross-resistance by mosquito P450s that metabolise pyrethroids. Sci Rep. 2020;10(1):16232.

66. Chandor-Proust A, Bibby J, Regent-Kloeckner M, Roux J, Guittard-Crilat E, Poupardin R, et al. The central role of mosquito cytochrome P450 CYP6Zs in insecticide detoxification revealed by functional expression and structural modelling. Biochem J. 2013;455(1):75–85.

67. Antonio-Nkondjio C, Poupardin R, Tene BF, Kopya E, Costantini C, Awono-Ambene P, et al. Investigation of mechanisms of bendiocarb resistance in Anopheles gambiae populations from the city of Yaounde, Cameroon. Malar J. 2016;15(1):424.

68. Riveron JM, Ibrahim SS, Mulamba C, Djouaka R, Irving H, Wondji MJ, et al. Genome-Wide Transcription and Functional Analyses Reveal Heterogeneous Molecular Mechanisms Driving Pyrethroids Resistance in the Major Malaria Vector Anopheles funestus Across Africa. G3 (Bethesda). 2017;7(6):1819–32.

69. Vontas J, Grigoraki L, Morgan J, Tsakireli D, Fuseini G, Segura L, et al. Rapid selection of a pyrethroid metabolic enzyme CYP9K1 by operational malaria control activities. Proceedings of the National Academy of Sciences of the United States of America. 2018;115(18):4619–24.

70. Hearn J, Tagne CD, Ibrahim SS, Tene-Fossog B, Mugenzi LJ, Irving H, et al. Multi-omics analysis identifies a CYP9K1 haplotype conferring pyrethroid resistance in the malaria vector Anopheles funestus in East Africa. bioRxiv 2022;2021.10.21.465247.

71. Kefi M, Charamis J, Balabanidou V, Ioannidis P, Ranson H, Ingham VA, et al. Transcriptomic analysis of resistance and short-term induction response to pyrethroids, in Anopheles coluzzii legs. BMC Genomics. 2021;22(1):891.

72. Dunse KM, Kaas, Q., Guarino, R.F., Barton, P.A., Craik, D.J. and Anderson, M.A. Molecular basis for the resistance of an insect chymotrypsin to a potato type II proteinase inhibitor. Proceedings of the National Academy of Sciences. 2010;107(34):15016–21.

73. Hu HX, Zhou D, Ma L, Shen B, Sun Y, Zhu CL. Lipase is associated with deltamethrin resistance in Culex pipiens pallens. Parasitol Res. 2020;119(1):23–30.

74. Weetman D, Wilding CS, Neafsey DE, Muller P, Ochomo E, Isaacs AT, et al. Candidate-gene based GWAS identifies reproducible DNA markers for metabolic pyrethroid resistance from standing genetic variation in East African Anopheles gambiae. Sci Rep. 2018;8(1):2920.

75. Weill M, Malcolm C, Chandre F, Mogensen K, Berthomieu A, Marquine M, et al. The unique mutation in ace-1 giving high insecticide resistance is easily detectable in mosquito vectors. Insect Mol Biol. 2004;13(1):1–7.

76. Grau-Bove X, Lucas E, Pipini D, Rippon E, van ’t Hof AE, Constant E, et al. Resistance to pirimiphos-methyl in West African Anopheles is spreading via duplication and introgression of the Ace1 locus. PLoS Genet. 2021;17(1):e1009253.

77. Williams J, Cowlishaw R, Sanou A, Ranson H, Grigoraki L. In vivo functional validation of the V402L voltage gated sodium channel mutation in the malaria vector An. gambiae. Pest Manag Sci. 2022;78(3):1155–63.

78. Clarkson CS, Miles A, Harding NJ, O’Reilly AO, Weetman D, Kwiatkowski D, et al. The genetic architecture of target-site resistance to pyrethroid insecticides in the African malaria vectors Anopheles gambiae and Anopheles coluzzii. Mol Ecol. 2021;30(21):5303–17.

79. Ding Y, Hawkes N, Meredith J, Eggleston P, Hemingway J, Ranson H. Characterization of the promoters of Epsilon glutathione transferases in the mosquito Anopheles gambiae and their response to oxidative stress. Biochem J. 2005;387(Pt 3):879–88.

80. Balabanidou V, Kampouraki A, MacLean M, Blomquist GJ, Tittiger C, Juarez MP, et al. Cytochrome P450 associated with insecticide resistance catalyzes cuticular hydrocarbon production in Anopheles gambiae. Proceedings of the National Academy of Sciences of the United States of America. 2016;113(33):9268–73.

81. Vannini L, Augustine Dunn W, Reed TW, Willis JH. Changes in transcript abundance for cuticular proteins and other genes three hours after a blood meal in Anopheles gambiae. Insect biochemistry and molecular biology. 2014;44:33–43.

82. Messenger LA, Impoinvil LM, Derilus D, Yewhalaw D, Irish S, Lenhart A. A whole transcriptomic approach provides novel insights into the molecular basis of organophosphate and pyrethroid resistance in Anopheles arabiensis from Ethiopia. Insect biochemistry and molecular biology. 2021;139:103655.

83. Zhao LaJ, W.A. . Expression of heat shock protein genes in insect stress responses. . Invertebrate Survival Journal. 2012;9(1):93–101.

84. Cassone BJ, Molloy MJ, Cheng C, Tan JC, Hahn MW, Besansky NJ. Divergent transcriptional response to thermal stress by Anopheles gambiae larvae carrying alternative arrangements of inversion 2La. Mol Ecol. 2011;20(12):2567–80.

85. VanHook AM. Understanding hygrosensation: How flies sense changes in humidity. Science Signaling. 2016;9(430):ec127-ec.

86. Enjin A, Zaharieva EE, Frank DD, Mansourian S, Suh GS, Gallio M, et al. Humidity Sensing in Drosophila. Curr Biol. 2016;26(10):1352–8.

87. Greppi C, Laursen WJ, Budelli G, Chang EC, Daniels AM, van Giesen L, et al. Mosquito heat seeking is driven by an ancestral cooling receptor. Science. 2020;367(6478):681–4.

