## Supplementary Text - Methods and Results for "Molecular drivers of insecticide resistance in the Sahelo-Sudanian populations of a major malaria vector *Anopheles coluzzii*"

***Cloning of GSTe2 and CYP6Z2 5’ regulatory element into PGL3-Basic vector and dual luciferase reporter assay***

The *An. gambiae* cell line 4a-3B (MRA-919, https://www.beiresources.org/) were maintained at 25 °C in Schneider’s insect medium (SIGMA, MO, USA) supplemented with 10 % (v/v) heat inactivated foetal bovine serum (FBS) (SIGMA) and 1% penicillin/streptomycin (SIGMA). Approximately 4 × 10^4^ of cells per well were plated out 24 h before transfection into 24-well plates and allowed to reach 60-70% confluence. At about ~70% confluence, constructs were transfected into the cells using Qiagen Effectene Transfection Reagent (QIAGEN, Hilden, Germany). These constructs include either 200 ng recombinant reporter constructs of *GSTe2* promoters, LRIM promoter in pGL3-Basic vector (Lynd and Lycett, 2011), or promoter-less pGL3-Basic control. The constructs were co-transfected together with 1 ng/µL of internal control, sea pansy *Renilla reniformis* luciferase containing the *Drosophila* Actin *5C* promoter in pRL-null (Lynd and Lycett, 2011). The constructs were diluted together with the *Renilla* plasmid in 50 µL DNA condensation buffer, followed by 1.6 µL enhancer, briefly vortexed for 1 s and incubated for 1.5 min at room temperature. Tubes were microfuged for 1 s to collect drops before 5 µL of Effectene transfection reagent was added with pipetting up and down 5 times. Tubes were incubated for 7.5 min at room temperature to allow formation of transfection complex before 350 µL of growth medium was added with mixing. Plated out cells were washed with 3 mL PBS and 350 µL fresh growth medium containing FBS and antibiotics added. The transfection complex was added to the cells in the plates dropwise with gentle swirling to mix. For each experiment transfection was done in triplicates for each construct. Transfected cells were incubated at 25 °C for 48 h to allow protein expression before the cells were washed with PBS and lysed in 100 ml of 1x passive lysis buffer (Promega). The activities of the firefly luciferase were measured with a luminometer (EG & G Berthold, Baden-Württemberg, Germany), using a Dual-Luciferase Reporter Assay kit (Promega) with normalisation using the *Renilla* luciferase activity*.* Protocol for cell lysis and reporter assay was as outlined in the Promega Quick Protocol (https://www.promega.com/-/media/files/resources/protcards/dual-luciferase-reporter-assay-and-dual-luciferase-reporter-1000-assay-systems-quick-protocol.pdf).

***Cloning and microinjection of GSTe2 and CYP6Z2 in Drosophila melanogaster flies***

Amplification of full-length *GSTe2* and *CYP6Z2* was carried out using Phusion High-Fidelity DNA Polymerase, with *trg* primers bearing *Bgl*II and *Xba*I (Table S4). PCR products were cleaned and cloned into the pUASattB vector linearised with the above restriction enzymes. Using the PhiC31 system, clones were injected into the germ-line of *D. melanogaster* line carrying the attP40 docking site, 25C6 on chromosome 2 [y w M (eGFP, vas-int, dmRFP) ZH-2A; P{CaryP}attP40]. Microinjection and balancing of UAS stock to remove integrase was carried out by the Fly Facility (Cambridge, UK) generating UAS-GSTe2 and UAS-CYP6Z2 transgenic lines. Ubiquitous expression of the transgene in adult F_1_ progeny (the experimental group) was attained following crossing of virgin females from the GAL4-Actin driver strain Act5C-GAL4, BL25374 [y[1] w[*]; P{Act5C-GAL4-w}E1/CyO, 1;2] (Bloomington, IN, USA) with male flies from the UAS-lines. For control group, adult F_1_ progeny with the same genetic background as the experimental group but without GSTe2 or CYP6Z2 insertion were obtained by crossing virgin females from the driver strain Act5C-GAL4 with the UAS-null recipient males with white eyes (devoid of pUASattB-GSTe2 or pUASattB-CYP6Z2 insertions).

**Results**

***Gene Ontology Enrichment Analysis***

For C vs S comparison (upregulated in C) the glutathione S-transferases (the most enriched term), oxidoreductase (frequency = 2.11 %), the monooxygenase (frequency = 1.21 %), hydrolase, chitin binding, and oxidant activities were over-represented (Figure S6e). This is in contrast with the terms downregulated in C, were ion binding and channelling, and neurotransmitter activities were the most over-represented (Figure S6f). ATP binding has the highest frequency (13.64 %), followed by zinc ion binding (3.63 %).

For the rest of the three countries contrasts were also observed. For example, comparison of R vs S (upregulated in R) in populations from Niger, Chad, and Cameroon, revealed glutathione S-transferase activities over-represented in Niger and Chad, C-C lyase, DNA replication origin binding and RNA cis-regulatory region binding activities in Niger, sulfotransferase and DNA and RNA binding activities in Chad, as well as NADH dehydrogenase activities in Cameroon (Figure S6g, -n and -u). The terms down-regulated in R include ion transport and channelling, and transmembrane transport, as well as peptidase activities in Niger and Chad, while carbohydrate transporter, ligase, antioxidant, alkene-1-monoxygenase and disulfide oxidoreductase activities were over-represented in Cameroon (Figure S6h, -p and -v).

For comparison of R vs C (upregulated in R), the common over-represented terms include oxidoreductase activities in all the three countries, chitin binding and transport activities in Niger and Chad, peroxidase in Niger, and acyl transferase activities in Cameroon (Figure S6j, -q and -w); while terms down-regulated in R include RNA transcription and DNA binding factors activities were over-represented in Niger and Chad (Figure S6k, -r and -x).

For comparison of C vs S (upregulated in C), glutathione S-transferase and oxidoreductase activities were over-represented in all the three countries; peroxidase activities were over-represented in Niger and Chad, nucleoside triphosphate activities in Niger and Cameroon, chitin binding in Chad, and carbohydrate transporter activities in Niger and Cameroon (Figure S6l, -s and -y). The terms downregulated in C include ion binding and channelling, neurotransmission and transmembrane transporter activities in Niger, DNA and RNA binding, dynein chain binding and structural constituents of ribosome activities in Chad, peptidase, phosphatase, ATPase, lipase and protein binding in Cameroon (Figure S6m, -t and z).

***Detection of Signatures of Selective Sweeps***

Three heat shock proteins, *hsp70* (1/8, AGAP004944), *hsp83* (AGAP006958) and *hsp110* (AGAP010331) were possibly undergoing selection as well. For example, the *hsp83* (Tajima’s D = -1.48 for Chad, -1.44 for Nigeria and -1.09 for Niger, but with combined *F*st of 0.17); *hsp110* (Tajima’s D = -1.03 in Chad and -0.85 in Nigeria, combined *F*st = 0.29). Positive selections were also evident in the *GSTe1* (AGAP009195, Tajima’s D = -1.65 in Chad and -1.81 in Niger, *F*st = 0.03) and *GSTU1* (AGAP000947, Tajima’s D = -1.46 in Nigeria and -0.98 in Cameroon, *F*st = 0.44). Surprisingly, three GST genes, *GSTe3*, *GSTe4* and *GSTe7* were under selection in both the field populations, and the Ngoussou, while other genes were undergoing selection only in the Ngoussou [e.g., an alkaline phosphatase (AGAP011302), an *acetylcholinesterase*-2 (AGAP000466), *CY4G17*, and *GSTD3*]. Other genes undergoing selection, but not in the list of top 100 metabolic resistance genes were provided in the File S3 (sheet 1, rows 21-40).

LYND, A. & LYCETT, G. J. 2011. Optimization of the Gal4-UAS system in an Anopheles gambiae cell line. *Insect Mol Biol,* 20**,** 599-608.
