## Supplementary Tables for "Molecular drivers of insecticide resistance in the Sahelo-Sudanian populations of a major malaria vector *Anopheles coluzzii*"

**Additional Tables**

**Table S1: RNAseq descriptive statistics from flagstat output files: pre-alignment statistics**

| **Sample name** | **Untrimmed reads** | **Trimmed reads** | **R1/R2 pairs ^1^** | **R0 reads (%) ^2^** | **Salmon aligned pairs^3^ (%)** |
| --- | --- | --- | --- | --- | --- |
| NGUSSO1 | 23236022 | 23085050 | 11467688 | 149674 (0.65) | 8685780 (76) |
| NGUSSO2 | 21846456 | 21683616 | 10760797 | 162022 (0.75) | 8241274 (77) |
| NGUSSO3 | 36064202 | 35834909 | 17803580 | 227749 (0.66) | 12745304 (72) |
| TAKDELTA1 | 38357800 | 38105880 | 18927771 | 250338 (0.66) | 13654716 (72) |
| TAKDELTA3 | 37422996 | 37155425 | 18444847 | 265731 (0.72) | 13752204 (75) |
| TAKDELTA5 | 23224936 | 23100393 | 11488482 | 123429 (0.53) | 8303633 (73) |
| TAKUNEXPRNA1 | 40406142 | 40159798 | 19957677 | 244444 (0.61) | 15367095 (77) |
| TAKUNEXPRNA4 | 36419154 | 36139911 | 17931231 | 277449 (0.77) | 13367573 (75) |
| TAKUNEXPRNA6 | 34912884 | 34718932 | 17263395 | 192142 (0.55) | 13588558 (79) |
| SIMATDELTA1 | 21200894 | 21043894 | 10444002 | 155890 (0.74) | 7351634 (70) |
| SIMATDELTA2 | 19074810 | 18932389 | 9395432 | 141525 (0.48) | 6650890 (71) |
| SIMATDELTA3 | 29720372 | 29557954 | 14698449 | 161056 (0.54) | 10440312 (71) |
| SIMATUNEXP2 | 32672736 | 32483686 | 16147908 | 187870 (0.58) | 11780452 (73) |
| SIMATUNEXP3 | 5866494 | 5838421 | 2905228 | 27965 (0.48) | 2394984 (82) |
| SIMATUNEXP4 | 40935832 | 40615430 | 20148507 | 318416 (0.78) | 14422724 (72) |
| HADRES3RNA | 36551724 | 36298866 | 18023985 | 250896 (0.69) | 13605875 (75) |
| HADRES5RNA | 27971672 | 27795464 | 13810284 | 174896 (0.63) | 10533382 (76) |
| HADRES6RNA | 23774116 | 23541492 | 11655070 | 231352 (0.98) | 8949568 (77) |
| HADUNEXP1RNA | 28787580 | 28621210 | 14227987 | 165236 (0.58) | 10758132 (76) |
| HADUNEXP3RNA | 35595388 | 35314457 | 17517601 | 279255 (0.79) | 13241662 (76) |
| HADUNEXP6RNA | 27478724 | 27340671 | 13601827 | 137017 (0.50) | 10199851 (75) |
| [CHAD_DELTA_1_RNA](http://cgr.liv.ac.uk/illum/LIMS18211_0f587c04e76299c6/Raw/Sample_1-CHAD_DELTA_1_RNA) | 39348916 | 39015513 | 19341653 | 332207 (0.85) | 14718325 (76) |
| [CHAD_DELTA_2_RNA](http://cgr.liv.ac.uk/illum/LIMS18211_0f587c04e76299c6/Raw/Sample_2-CHAD_DELTA_2_RNA) | 38979368 | 38642265 | 19153149 | 335967 (0.87) | 14098280 (74) |
| [CHAD_DELTA_3_RNA](http://cgr.liv.ac.uk/illum/LIMS18211_0f587c04e76299c6/Raw/Sample_3-CHAD_DELTA_3_RNA) | 44559028 | 44089639 | 21810778 | 468083 (1.06) | 16253197 (75) |
| [CHAD_UNX_1_RNA](http://cgr.liv.ac.uk/illum/LIMS18211_0f587c04e76299c6/Raw/Sample_4-CHAD_UNX_1_RNA) | 36888528 | 36447976 | 18004243 | 439490 (1.21) | 13621400 (76) |
| [CHAD_UNX_2_RNA](http://cgr.liv.ac.uk/illum/LIMS18211_0f587c04e76299c6/Raw/Sample_5-CHAD_UNX_2_RNA) | 44924880 | 44478094 | 22016328 | 445438 (1.00) | 16664846 (76) |
| [CHAD_UNX_3_RNA](http://cgr.liv.ac.uk/illum/LIMS18211_0f587c04e76299c6/Raw/Sample_6-CHAD_UNX_3_RNA) | 37484660 | 37100903 | 18359186 | 382531 (1.03) | 13724852 (75) |

^1^ Forward (R1) and reverse (R2) read pairs after trimming; ^2^ Reads unpaired after trimming (% of total trimmed reads); ^3^ Reads aligned to *An. gambiae* transcriptome (AgamP4.10) by Salmon. NGUSSO = NGOUSSOU (THE FULLY SUSCEPTIBLE ANOPHELES COLUZZII COLONY, ORIGINATED FROM CAMEROON); TAK = TAKATSABA, NIGER REPUBLIC; SIM = SIMATOU CAMEROON; HAD = HADIYAU, NIGERIA; CHAD = CHAD REPUBLIC; UNEXP AND UNX = UNEXPOSED.

**Table S2: RNAseq descriptive statistics from flagstat output files: post-alignment statistics**

| **Sample ID** | **Reads to align (R1+R2)** | **Aligned reads (%) ^1^** | | | **Aligned reads, filtered (%) ^1,2^** | **Aligned in pair (%) ^3^** | **Singleton (%) ^3^** |
| --- | --- | --- | --- | --- | --- | --- | --- |
| NGUSSO1 | 22935376 | 21380943 (93.22) | | 21380943 (93.22) | | 21061238 (98.50) | 319705 (1.50) |
| NGUSSO2 | 21521594 | 20186461 (93.8) | | 20186461 (93.8) | | 19899040 (98.58) | 287421 (1.42) |
| NGUSSO3 | 35607160 | 32568174 (91.47) | | 32568174 (91.47) | | 32054416 (98.42) | 513758 (1.58) |
| TAKDELTA1 | 37855542 | 37855542 (93.03) | | 35216388 (93.01) | | 34509477 (97.99) | 706911 (2.01) |
| TAKDELTA3 | 36889694 | | 34517594 (93.57) | 34517594 (93.57) | | 33879690 (98.15) | 637904 (1.85) |
| TAKDELTA5 | 22976964 | | 21540030 (93.75) | 21540030 (93.75) | | 21144488 (98.16) | 395542 (1.84) |
| TAKUNEXPRNA1 | 39915354 | | 37583736 (94.16) | 37583736 (94.16) | | 36931604 (98.26) | 652132 (1.74) |
| TAKUNEXPRNA4 | 35862462 | | 33708881 (93.99) | 33708878 (93.99) | | 33091957 (98.17) | 616921 (1.83) |
| TAKUNEXPRNA6 | 34526790 | | 32368261 (93.75) | 32368261 (93.75) | | 31816296 (98.29) | 551965 (1.71) |
| SIMATDELTA1 | 20888004 | | 19342520 (92.60) | 19342519 (92.60) | | 18967519 (98.06) | 375000 (1.94) |
| SIMATDELTA2 | 18790864 | | 17289997 (92.01) | 17289997 (92.01) | | 16934904 (97.95) | 355093 (2.05) |
| SIMATDELTA3 | 29396898 | | 27331837 (92.98) | 27331836 (92.98) | | 26790118 (98.02) | 541718 (1.98) |
| SIMATUNEXP2 | 32295816 | | 29918068 (92.64) | 29918068 (92.64) | | 29360698 (99.14) | 557370 (1.68) |
| SIMATUNEXP3 | 5810456 | | 5322424 (91.60) | 5322424 (91.60) | | 5251426 (98.67) | 70998 (1.33) |
| SIMATUNEXP4 | 40297014 | | 36118764 (89.63) | 36118762 (89.63) | | 35397250 (98.00) | 721512 (1.43) |
| HADRES3RNA | 36047970 | | 33713459 (93.52) | 33713458 (93.52) | | 33107926 (98.20) | 605532 (1.80) |
| HADRES5RNA | 27620568 | | 25899936 (93.77) | 25899936 (93.77) | | 25448202 (98.26) | 451734 (1.74) |
| HADRES6RNA | 23310140 | | 21712521 (93.15) | 21712521 (93.15) | | 21320452 (98.19) | 392069 (1.81) |
| HADUNEXP1RNA | 28455974 | | 26719962 (93.90) | 26719962 (93.90) | | 26293680 (98.40) | 426282 (1.60) |
| HADUNEXP3RNA | 35035202 | | 32884162 (96.86) | 32884162 (96.86) | | 32321622 (98.29) | 562540 (1.71) |
| HADUNEXP6RNA | 27203654 | | 25441368 (93.52) | 25441365 (93.52) | | 25045653 (98.44) | 395712 (1.56) |
| [CHAD_DELTA_1_RNA](http://cgr.liv.ac.uk/illum/LIMS18211_0f587c04e76299c6/Raw/Sample_1-CHAD_DELTA_1_RNA) | 38683306 | | 36184989 (93.54) | 36184988 (93.54) | | 35497570 (98.10) | 687418 (1.90) |
| [CHAD_DELTA_2_RNA](http://cgr.liv.ac.uk/illum/LIMS18211_0f587c04e76299c6/Raw/Sample_2-CHAD_DELTA_2_RNA) | 38306298 | | 35787249 (93.42) | 35787249 (93.42) | | 35108908 (98.10) | 678341 (1.90) |
| [CHAD_DELTA_3_RNA](http://cgr.liv.ac.uk/illum/LIMS18211_0f587c04e76299c6/Raw/Sample_3-CHAD_DELTA_3_RNA) | 43621556 | | 40741028 (93.40) | 40741027 (93.40) | | 39774810 (97.63) | 966217 (2.37) |
| [CHAD_UNX_1_RNA](http://cgr.liv.ac.uk/illum/LIMS18211_0f587c04e76299c6/Raw/Sample_4-CHAD_UNX_1_RNA) | 36008486 | | 33680160 (93.53) | 33680159 (93.53) | | 32989999 (97.95) | 690160 (2.05) |
| [CHAD_UNX_2_RNA](http://cgr.liv.ac.uk/illum/LIMS18211_0f587c04e76299c6/Raw/Sample_5-CHAD_UNX_2_RNA) | 44032656 | | 41186880 (93.54) | 41186879 (93.54) | | 40374965 (98.03) | 811914 (1.97) |
| [CHAD_UNX_3_RNA](http://cgr.liv.ac.uk/illum/LIMS18211_0f587c04e76299c6/Raw/Sample_6-CHAD_UNX_3_RNA) | 36718372 | | 34369490 (93.60) | 34369490 (93.60) | | 33677582 (97.99) | 691908 (2.01) |
| GOUNPERM2RNA | 24924672 | | 23402379 (93.89) | 23402379 (93.89) | | 23002742 (98.29) | 399637 (1.71) |
| GOUNPERM4RNA | 24507610 | | 23055920 (94.08) | 23055920 (94.08) | | 22683092 (98.38) | 372828 (1.62) |
| GOUNPERM7RNA | 29030956 | | 27275670 (93.95) | 27275666 (93.95) | | 26831596 (98.37) | 444070 (1.63) |
| GOUNUNEXP3RNA | 25156554 | | 23665448 (94.07) | 23665447 (94.07) | | 23299917 (98.46) | 365530 (1.54) |
| GOUNUNEXP4RNA | 24938738 | | 23091144 (92.59) | 23091144 (92.59) | | 22720240 (98.39) | 370904 (1.61) |
| GOUNUNEXP5RNA | 21633948 | | 20039292 (92.63) | 20039292 (92.63) | | 19723450 (98.42) | 315842 (1.58) |

^1^ % of reads to align; ^2^ Aligned reads filtered to remove reads with mapping quality <10; ^3^ % of filtered aligned reads with both read and its mate mapped to opposing strands of the reference sequence, with 3' ends innermost and 5' ends within the allowed distance from each other (0-500 bp).

**Table S3: List of primers used for qRT-PCR validation of the overexpressed genes across the Sahel**

| **Gene** | **Forward Primer** | **Reverse Primer** | **Amplicon Size (bp)** |
| --- | --- | --- | --- |
| *GSTe2* (AGAP009194) | ACCATTAATCTGCTAACGGGTG | AATTTACACGGGCCTGCTTG | 196 |
| *GSTZ1* (AGAP002898) | CAGCACTGCAACGAGTACC | CTTCTTCTCCTCACCGACGT | 249 |
| *CYP6Z2* (AGAP008218) | AGGCCACGAAGAACTACGAT | ACTTTTGCAGGAGTTGTGGC | 262 |
| *CYP4C27* (AGAP009246) | GGGTGTGAAGGTGAATGCTC | TTTCTTTGCCGACTCGAGTG | 232 |
| *CYP4G16* (AGAP001076) | TTTATCACGACGCCACATGC | AGCAATTCTATATCGCGTGGA | 213 |
| *CYP4G17* (AGAP000877) | TGTCACGACTACATGAGCGA | CGCAGGTGGATCTTCAGTTG | 158 |
| *CYP6P3* (AGAP002865) | AGCGGCTGAGAGAGGAAATT | GCTTCGGGATCACATGCTTT | 195 |
| *CYP6M2* (AGAP008212) | AGGTCGTGAGTGTGTGAGAG | CTTTCGAAGCCACACGGAAA | 152 |
| *CYP6Z3* (AGAP008217) | GCAAGGGAACACAGGTGATC | ACAAGCCCGATTTTGGACAC | 202 |
| *CYP9K1* (AGAP000818) | TCGCAGAAGCGGTCGGTTG | GAGTTCGTGCGCCATAAATGCAG | 117 |
| *UGT-B19* (AGAP007920) | TGGGAAAACGAAACGCTACC | CTTCGTAGGCCAGTACCACA | 241 |
| *COEBE3C* (AGAP005372) | CGGAAGCGATTCGAAACCAT | TCGTCCCGATGTGATAGAGC | 211 |
| *RPS7* (AGAP010592) | GTGTTCGGTTCCAAGGTGAT | TCCGAGTTCATTTCCAGCTC | 98 |
| *GPDH* (AGAP007593) | CTGCAAAAAGTCGATACCGC | CCTCGTACACGTACATCGTGA | 170 |

**Table S4: Primers used for the functional characterisation of candidate metabolic resistance genes**

| **Gene** | **Forward Primer** | | | | | **Reverse Primer** | |
| --- | --- | --- | --- | --- | --- | --- | --- |
| **Primers used for amplification of full-length cDNA sequences for *GSTe2* and *CYP6Z2*** | | | | | | | |
| ***GSTe2_Full*** | ATGTCCAACCTTGTACTGTAC | | | | | TTAAGCCTTAGCATTCTCCTC | |
| ***CYP6Z2_Full*** | ATGTTTGTTTACACTCTCGC | | | | | TCACTTTCTATGGTCTATCCTC | |
| ***Primers used for characterisation of 5’UTR (cis-regulatory elements) of GSTe2 and CYP6Z2*** | | | | | | | |
| ***3′-UTR_GSTe1_5′-UTR_GSTe2*** | CAGGTTCATGCTATTCGCGGTT | | | | | TTTCGCAGCGAACTAAAAACTGG | |
| ***3′-UTR_CYP6Z1_5′-UTR_CYP6Z2*** | GCAAAATACGCTGTGATAATTAGG | | | | | TATGCCACTATGCGTTTAGCC | |
| ***3′-UTR_CYP6Z1_5′UTR_CYP6Z2pGL3Basic*** | | GGTACCGCAAAATACGCTGTGATAATTAGG | | | | | AGATCTTATGCCACTATGCGTTTAGCC |
| ***3′-UTRGSTe1_5′-UTRGSTe2pGL3Basic*** | | | GGTACCCAGGTTCATGCTATTCGCGGTT | | | AGATCTTTTCGCAGCGAACTAAAAACTGG | |
| ***pGL3-Basic*** | | | ACTAGCAAAATAGGCTGTCCCCA | | | TGTTTTTGGCGTCTTCCATGGTG | |
| ***-308_GSTe2*** | | | GGTACCGCTAGCAAGTGTAGGATTTAAAGCACC | | |  | |
| ***-270_GSTe2*** | GGTACCGCTAGCAGTAACGTTTTTCTTGTGCAT | | |  | | | |
| ***-262_GSTe2*** | GGTACCGCTAGCGGAATTCGCTTCTGCTTTTATG | | | |  | | |
| ***-232_GSTe2*** | GGTACCGCTAGCCGAAGAACCGGAATGAGTATG | | |  | | | |
| ***GSTe2_minus_5’-UTR*** |  | | | CTCGAGAGATCTTTCAAGAGACACACCCGAACT | | | |
| **Primers used for *in vivo* functional characterization of *GSTe2* and *CYP6Z2* with transgenic analysis and flies qRT-PCR** | | | | | | | |
| ***trg_GSTe2*** | GAATTCATGTCCAACCTTGTACTGTAC | | | TCTAGATTAAGCCTTAGCATTCTCCTC | | | |
| ***trg_CYP6Z2*** | AGATCTATGTTTGTTTACACTCTCGC | | | TCTAGATCACTTTCTATGGTCTATCCTC | | | |
| ***qtrg_GSTe2*** | ACCATTAATCTGCTAACGGGTG | | | AATTTACACGGGCCTGCTTG | | | |
| ***qtrg_CYP6Z2*** | ACAATCCAGAAGCGATGGCAAA | | | CCTTCACGCACAAATCCAAGTA | | | |
| ***Dmel_RPL11*** | CGATCCCTCCATCGGTATCT | | | AACCACTTCATGGCATCCTC | | | |

Bright green is *kpn*I, pink is *BgL*II*,* teal is *EcoR*I, yellow is *Nhe*I, turquoise = *Xho*I, red = *Xba*I, dark red = *BamH*I, dark green = RAKRR coding for furin proteinase cleavage sequences, italics and underlined = the abolished *CYP6Z2* stop codon substituted for arginine, grey = *Nde*I.

**Table S5: Ploidy scores and frequencies of chromosomal inversion polymorphisms**

| **Ploidy Score (Frequency)** | | | | | | |
| --- | --- | --- | --- | --- | --- | --- |
| Population Inversions | | | | | | |
|  | 2La | 2Rb | 2Rc | 2Rd | 2Rj | 2Ru |
| Nigeria | 16.000 (0.100) | 12.641 (0.790) | 14.166 (0.885) | 0.292 (0.018) | 1.823 (0.303) | 1.025 (0.064) |
| Niger | 15.986 (0.999) | 13.576 (0.848) | 13.683 (0.855) | 0.476 (0.0297) | 1.161 (0.0726) | 0.588 (0.0367) |
| Chad | 15.873 (0.992) | 12.836 (0.802) | 14.211 (0.888) | 0.477 (0.0298) | 1.077 (0.0673) | 0.127 (0.0079) |
| Cameroon | 16 (0.100) | 4.650 (0.2908) | 7.518 (0.469) | 0.799 (0.049) | 0.523 (0.032) | 3.887 (0.242) |
| Ngoussou | 1.001 (0.062) | 0.855 (0.0534) | 1.583 (0.0989) | 0.044 (0.002) | 2.091 (0.130) | 0.00 |

Ploidy score = average score for all replicates from the same population; frequency is the ratio of the ploidy score in relation to the total of the possible ploidy (16).

**Table S6: Summary statistics for polymorphisms of *GSTe2* 5’-UTR from Sahel countries.**

| **Population/Phenotype** | **n** | **S** | **Syn** | **Nonsyn** | **h** | **H_d_** | **π (k)** | **D (Tajima)** | **D* (Fu and Li)** |
| --- | --- | --- | --- | --- | --- | --- | --- | --- | --- |
| **Nigeria-alive** | 10 | 16 | 16 | 0 | 4 | 0.711 | 0.0208 | 1.373 ^ns^ | 0.799 ^ns^ |
| **Nigeria-dead** | 10 | 1 | 1 | 0 | 2 | 0.200 | 0.00057 | -1.1117 ^ns^ | -1.346 ^ns^ |
| **Niger-alive** | 10 | 3 | 3 | 0 | 2 | 0.533 | 0.00456 | 1.8305 ^ns^ | 1.154 ^ns^ |
| **Niger-dead** | 10 | 0 | 0 | 0 | 1 | 0.000 | 0.000 | 0.000 | 0.000 |
| **Chad-alive** | 10 | 13 | 13 | 0 | 2 | 0.356 | 0.01317 | 0.0266 ^ns^ | 1.499 ^ns^ |
| **Chad-dead** | 10 | 0 | 0 | 0 | 1 | 0.000 | 0.000 | 0.000 | 0.000 |
| **Cameroon-alive** | 10 | 13 | 13 | 0 | 2 | 0.467 | 0.01728 | -1.456 ^ns^ | 1.499 ^s^ |
| **Cameroon-dead** | 10 | 0 | 0 | 0 | 1 | 0.000 | 0.000 | 0.000 | 0.000 |
| **Ngoussou** | 10 | 0 | 0 | 0 | 1 | 0.000 | 0.000 | 0.000 | 0.000 |
| **All** | 90 | 17 | 17 | 0 | 6 | 0.299 | 0.0074 | -0.635 ^ns^ | -0.297 ^ns^ |

n = number of sequences (n); S, number of polymorphic sites; h, haplotype; H_d_, haplotype diversity Syn, Synonymous mutations; Nonsyn, Non-synonymous mutations; π, nucleotide diversity (k= mean number of nucleotide differences); Tajima’s D and Fu and Li’s D statistics, ns, not significant, s, significant at p <0.02.
